# Sexually dimorphic interzonal crosstalk reshapes the adrenal cortex in response to pathophysiological challenges

**DOI:** 10.1101/2025.09.05.674435

**Authors:** Nicolo Faedda, Alaa B Abdellatif, Francesco Carbone, May Fayad, Bakhta Fedlaoui, Rita Chamoun, Romane L’Héritier, Ilaria Del Gaudio, Ourida Koual, Isabelle Giscos-Douriez, Stéphanie Baron, Ben Atkinson, Xiaomin Li, Linghui Kong, Yunling Xu, Eric Camerer, Mickael Menager, Fabio L. Fernandes-Rosa, Sheerazed Boulkroun, Maria-Christina Zennaro

## Abstract

**Structured Abstract:** 

**Background:** Primary aldosteronism is the most common form of secondary arterial hypertension, due to autonomous aldosterone production from the adrenal cortex. Genome wide association studies discovered genetic risk loci associated with the disease, which may affect adrenal cortex renewal, differentiation and lineage conversion. Genetic susceptibility may be modulated by environmental challenges.

**Method:** Here we investigate how environmental cues affecting mineralocorticoid output modulate adrenal cortex homeostasis, cell lineage conversion and zone-specific transcriptional landscape. We have used a newly developed *Cyp11b2^Cre^*mouse model and characterised its adaptation to a high or low salt diet (HSD, LSD), as well as dexamethasone (DEX) treatment. To explore underlying mechanisms, deep functional and morphological phenotyping, lineage tracing and spatial transcriptomics of the adrenal cortex were performed.

**Results:** *Cyp11b2* expression was detected in *Cyp11b2^Cre^*-*m*T*m*G mice as early as day P1 in different areas of the ZG, associated with high plasma aldosterone levels, with lineage conversion of zona glomerulosa (ZG) into zona fasciculata (ZF) cells progressing between 2 and 9 weeks of age and a progressive reduction of ZG size and evolution of cell components of the adrenal cortex over time. Transdifferentiation progressed into the X-zone (ZX) in females, revealing a previously unrecognized connection between ZF and ZX cells in adult mice. A sexually dimorphic, reciprocal interaction between the ZG and the ZF in adapting to salt diets or DEX treatment was observed, involving changes in cell composition and transcriptional reprogramming of the three zones.

**Conclusion:** HSD, LSD and DEX induce a sexually dimorphic cellular and transcriptional response, involving all layers of the adrenal cortex, and the reciprocal contribution of ZG and ZF cells, indicating functional interaction between adrenocortical zones in adapting to external cues.

## Introduction

Primary aldosteronism (PA) is the most common form of secondary arterial hypertension, affecting up to 20% of hypertensive patients and contributing to increased cardiovascular risk^1–3^. It is due to excessive and autonomous aldosterone production by the adrenal gland, due in the majority of cases to a unilateral aldosterone producing adenoma (APA) or bilateral adrenal hyperplasia (BAH)^4^. Recent evidence suggests that abnormalities in aldosterone production may contribute to a significant fraction of hypertension in the general population^5^. While several germline and somatic mutations have been described in familial forms of PA and APA^5–7^, recent genome wide association studies (GWAS) discovered genetic loci associated with increased risk of developing the disease^8–10^. These loci are shared between APA and BAH, revealing a hitherto unexpected common pathophysiology between unilateral and bilateral PA. Remarkably, PA risk loci also overlap with risk loci for hypertension traits, including resistant hypertension, providing genetic support to the clinical evidence of a pathophysiological continuum of aldosterone dysregulation throughout the spectrum of blood pressure, from normotension to PA^5^. Mechanistic studies have shown that genes located within risk loci for PA identified in GWAS modify adrenocortical function in cell and mouse models^8,11^. Experimental data on *CASZ1* and *RXFP2*, two PA susceptibility genes, suggest a potential pathogenic mechanism whereby a genetically determined reduction in aldosterone production by the ZG in carriers of PA risk alleles leads to PA due to lifelong increased stimulation of the adrenal cortex to maintain adequate mineralocorticoid output^8^. This chronic stimulatory drive may promote increased cell proliferation in the ZG, eventually resulting in a propitious environment for the appearance of somatic mutations within single or multiple aldosterone-producing areas in one or both adrenal glands^8^. On the other hand, *Wnt2b*, another PA susceptibility gene, is expressed in capsular cells of the adrenal cortex. Its inactivation in mice led to dysmorphic and hypocellular ZG, with impaired aldosterone production^11^. It therefore appears that risk loci for PA mainly affect adrenal cortex maintenance and cell differentiation, rather than directly aldosterone production. This raises the hypothesis that common genetic variation may modulate adrenal cortex renewal, differentiation and cell lineage conversion, eventually leading to hyperplasia, nodulation and PA in extreme cases.

The adrenal cortex consists of different concentric cell layers with different functional properties. The outermost layer, the zona glomerulosa (ZG), plays a key role in the renin-angiotensin-aldosterone system (RAAS) by producing mineralocorticoids, primarily aldosterone, which is essential for regulating salt and fluid balance in the kidney. The middle layer, the zona fasciculata (ZF), produces glucocorticoids, cortisol in humans and corticosterone in mice, under the influence of the hypothalamic-pituitary-adrenal (HPA) axis, affecting glucose homeostasis and the body’s chronic stress response. The innermost layer, the zona reticularis (ZR) in humans, is responsible for androgen production. In mice, this layer is absent; instead, an X-zone (ZX) derived from the foetal adrenal cortex is present, which has an unknown function and regresses at puberty in males and after the first pregnancy in females^12–15^. This so-called “functional” zonation of the adrenal cortex results from the zone-specific expression of steroidogenic enzymes, including aldosterone synthase (AS, encoded by *CYP11B2*) in the ZG and 11β-hydroxylase (encoded by *CYP11B1*) in the ZF. The adrenal cortex is a highly dynamic and adaptive tissue, continuously renewing in adult mice and able to adapt its function and size to environmental requirements, with important sex-specific differences^16^. Renewal takes place in the outer layers of the adrenal cortex, with newly formed cells undergoing centripetal migration, differentiation, and lineage conversion to establish the distinct functional steroidogenic zones^17–19^. Undifferentiated progenitor cells located in the subcapsular region develop into differentiated ZG cells, which subsequently transdifferentiate into ZF cells, although part of these progenitor cells may differentiate directly into ZF cells^18,19^. In mice, the ZX is considered to represent remnant tissue from the foetal cortex; lineage-tracing experiments have shown that the adult cortex derives from precursor cells in the fetal cortex, establishing a direct link between the adrenocortical zones^12–14^. Transdifferentiation has been described as the primary mechanism driving adult adrenal cortex renewal under physiological conditions, although the genes and pathways orchestrating these structural and metabolic changes remain a matter of investigation.

Genetic susceptibility may be modulated by environmental challenges. In particular, the adrenal cortex is constantly responding to pathophysiological demands with adaptations in hormone output and cellular plasticity^15,20,21^. Renewal and functional regeneration from glucocorticoid induced adrenocortical atrophy results from coordinated interplay between endocrine and paracrine regulation of growth and function involving different cell types and signalling pathways^22^. In the context of aldosterone homeostasis, salt diet is a major modulator of ZG homeostasis, leading to an adaptive hormonal and morphological response mediated mainly by the renin-angiotensin system^23,24^, but detailed changes in cell lineage conversion and zonal gene expression programs remain to be determined. The aim of this study was to explore the role of environmental cues in modulating adrenal cortex homeostasis, transdifferentiation and zone-specific transcriptional landscape, with a particular attention to the ZG and mineralocorticoid homeostasis, as well as sexual dimorphism. We used a newly developed *Cyp11b2*^Cre^ mouse model, in which aldosterone production is conserved, and investigated its adaptation to different salt diets as well as dexamethasone (DEX) treatment, by deep functional and molecular phenotyping of male and female mice. Characterisation of the new mouse model revealed the morphological and functional evolution of the adrenal cortex with age, with transdifferentiation progressing into the X-zone (ZX) in females, revealing a previously unrecognized connection between ZF and ZX cells in adult mice. We further discover a sexually dimorphic interaction between the ZG and the ZF in adapting to a high (HSD) or low salt diet (LSD) or DEX treatment through changes in cell composition and transcriptional reprogramming of the adrenal cortex.

## Results

### Generation and characterization of *Cyp11b2^Cre^* and *Cyp11b2^+/Cre^-mTmG* mice

We developed a novel *Cyp11b2^Cre^* mouse model in which the Cre recombinase is inserted within the *Cyp11b2* locus (exon 9) downstream of the stop codon located in the 3’ untranslated region (3’ UTR) of the gene, after an Internal Ribosome Entry Site (IRES) (Figure 1a). Insertion of the IRES allows the translation of two different proteins from a single mRNA transcript. The *Cyp11b2^+/Cre^* mice were crossed with *Gt(ROSA)26Sor^tm4(ACTB-tdTomato-EGFP)Luo/^J* reporter mice to generate the *Cyp11b2^Cre^*::*m*T*m*G line. Investigation of Cre recombination in this reporter model showed efficient and specific recombination in the ZG only, with no leakage in other tissues, such as brain, kidney, visceral and subcutaneous adipose tissue, aorta and mesenteric vessels (Figure S1a). Measurements of circulating hormones in plasma by liquid chromatography tandem mass spectrometry (LC-MS/MS), performed on 6-weeks old mice of both sexes carrying heterozygous (*Cyp11b2^+/Cre^)* or homozygous (*Cyp11b2^Cre/Cre^)* Cre alleles, showed normal plasma aldosterone levels compared to wild type, indicating normal aldosterone production (Figure 1b), as well as no major differences in overall corticosteroid profiles (Figure S1b). Plasma Renin Activity (PRA) showed no significant difference across genotypes, confirming a conserved RAAS (Figure 1b). While Western blot analyses showed a reduction of Aldosterone Synthase (AS) expression in *Cyp11b2^Cre/Cre^* mice compared to WT and *Cyp11b2^+/Cre^* mice in both sexes (Figure S1c), a similar number of AS*^+^* cells was detected by immunostaining across genotypes (Figure S1d). These results suggest that insertion of the Cre recombinase in the 3’ UTR of the *Cyp11b2* gene may have affected mRNA-stabilising sequences present in this region, possibly leading to faster transcript degradation and decreased protein levels; however, this reduction does not lead to any change in aldosterone production and renin levels. Nevertheless, to exclude any RAAS and ZG physiological variability across groups, only heterozygous mice were used for subsequent experiments.

**Figure 1.**
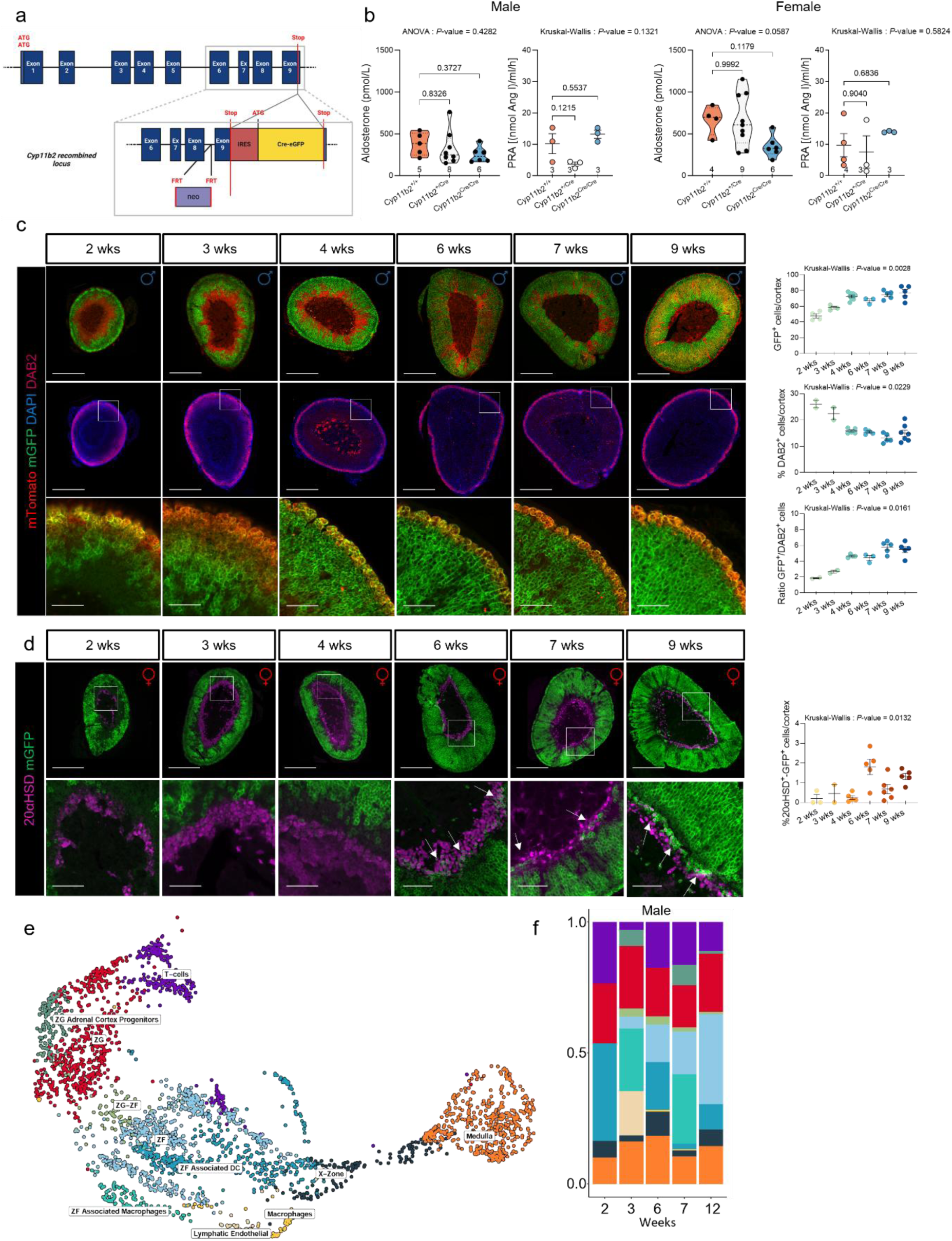
Characterization of the adrenal phenotype of the *Cyp11b2^Cre^* mouse model. **(a).** Cre recombinase was inserted into the 3’ untranslated region of the *Cyp11b2* gene, after an internal ribosomal entry site (IRES). The insertion of an IRES allows the translation of two different proteins from a single mRNA transcript, resulting in a functional aldosterone synthase protein. **(b).** Plasma aldosterone levels (ordinary one-way ANOVA, followed by Šídák’s multiple comparisons test) and Plasma Renin Activity (PRA, Kruskal-Wallis test followed by Dunn’s multiple comparisons test) of *Cyp11b2 ^+/+^* (orange), *Cyp11b2^+/Cre^* (white) and *Cyp11b2^Cre/Cre^* (light blue) male (left) and female (right) 6 weeks old mice. **(c).** Co-immunofluorescence of mGFP (green) with mTomato (red) and DAB2 (ruby red) of male *Cyp11b2^+/Cre^*-*m*T*m*G mouse adrenals at 2, 3, 4, 6, 7 and 9 weeks of age and quantification of mGFP^+^ cells and DAB2^+^ cells, with the analysis of cell lineage conversion expressed as the ratio of the number of cells expressing mGFP over the cortex area or the number of cells expressing DAB2. P values were obtained with Kruskal-Wallis test followed by Dunn’s multiple comparisons test. **(d).** Co-immunofluorescence of mGFP (green) and 20αHSD (purple) of female *Cyp11b2^+/Cre^*-*m*T*m*G mouse adrenals of 2, 3, 4, 6, 7 and 9 weeks of age. Quantification of 20αHSD positive cells and mGFP/20αHSD double-positive cells per cortex area (indicated by white arrows in the lower panel magnified image) at different time points. White squares represent the magnification area in the lower panel. The 7 weeks image has been used on Figure 8d. P values were obtained with Kruskal-Wallis test followed by Dunn’s multiple comparisons test. Scale Bar: 500 μm upper panels with entire adrenals and 100 μm lower panels. **(e).** uMAP of adrenal cell clusters from male *Cyp11b2^+/Cre^*-*m*T*m*G mice of 2, 3, 6, 7 and 12 weeks of age. **(f)**. Cell cluster proportions of male *Cyp11b2^+/Cre^*-*m*T*m*G mice of 2, 3, 6, 7 and 12 weeks of age. Data are shown as mean ± SEM. *P value < 0.05; **P value < 0.01; ***P value < 0.001.

### Lineage conversion progresses from ZF to ZX in *Cyp11b2^Cre^*::*m*T*m*G mice

The kinetics of adrenal cortex cell lineage conversion was investigated in male and female *Cyp11b2^Cre^*::*m*T*m*G mice from 2 weeks to 9 weeks of age by quantifying mGFP^+^ cells, which gradually replace mTomato^+^ cells as the cortex renewal process progresses^19^. During this time, adrenal weight increased progressively in both males and females (data not shown). At postnatal day 1 (P1), the surface of the adrenal showed large areas of mGFP^+^ cells (Figure S2a and S3a), whose localisation was restricted to the ZG (Figure S2a and S3b). At 2 weeks of age, the number of mGFP^+^ cells was 47.75±2.74% of adrenal cortex cells in males, increasing to 77.14±4.02% at 9 weeks of age (Figure 1c). In female mice, the mGFP^+^ cells similarly progressed from 41.54± 1.15% at 2 weeks to 75.12±3.93% at 9 weeks of age (Figure S2b). Comparing the evolution of mGFP^+^ cells in the cortex over time between sexes showed a significant difference at the age of 4 weeks, related to the major loss of ZX cells occurring at this age in males (Figure S2d). Quantification of Dab2^+^ cells, a marker of the ZG, showed a significant reduction of ZG size between 2 and 4 weeks of age in males, which was more progressive in females (Figure 1c and S2b). This was paralleled by a similar reduction of aldosterone levels over time (Figure S2c). Since the Cre recombinase is expressed under the control of the *Cyp11b2* promoter in the ZG, we also quantified the ratio of mGFP^+^ cells over Dab2^+^ cells, which showed a continuous increase between 2 and 7 weeks of age in both sexes (Figure 1c and S2b), with a slight but non significantly higher ratio after 6 weeks of age in females compared with males (Figure S2e). Adrenal 3D reconstruction of 6 weeks old mice by lightsheet microscopy revealed uneven penetration of mGFP^+^ cells in the adrenal cortex, suggesting spatially heterogenous transdifferentiation, as well as some columnar arrangements of mTomato cells in the ZF, connecting with the subcapsular region, in accordance with the alternative pathway of ZF differentiation moving directly from the subcapsular stem cell compartment to the ZF (Figure S1f). Remarkably, co-staining of mGFP and 20αHSD, a ZX marker (mGFP^+^/20αHSD^+^), was observed starting from 6 weeks in female *Cyp11b2^+/Cre^*-*m*T*m*G mice (Figure 1d), indicating a certain degree of lineage conversion between ZF and ZX.

Spatial transcriptomic (ST) analysis was performed on male and female *Cyp11b2^+/Cre^-m*T*m*G mice at 2, 3, 6, 7 and 12 weeks of age. Using well defined markers for the different cell populations composing the adrenal gland (Figure S4, S5 and Table S1), we identified 11 cell clusters (Figure 1e-f, S5a-c), including the ZG, ZF, ZX and medulla, as well as cluster containing both ZG and ZF cells (ZG-ZF cluster), a cluster containing ZG and adrenal cortex progenitors, T cells, macrophages, and ZF clusters enriched for dendritic cells (ZF associated DC) and macrophages (ZF associated macrophages), representing a mixture of ZG and immune cells. Partitioning the overall uMAP of male and female mice at different ages revealed that the process of adrenal growth was associated with the expansion or regression of different cell clusters composing the adrenal cortex (Figure 1f and S5d). In particular, ST confirmed the reduction over time of the ZG observed by immunolabeling, and revealed an evolution of the ZF during growth with changes of the 3 ZF containing clusters (pure ZF, ZF Associated DC and ZF Associated Macrophages) in both males and females.

These results indicate that *Cyp11b2* expression is detectable at P1 in different areas of the ZG, associated with a large Dab^+^ ZG as well as high aldosterone levels, both decreasing with age. Lineage conversion progresses into the ZX with time, becoming detectable around 6 weeks of age. These changes are associated with modifications in the cellular composition of the adrenal cortex.

### Low salt diet (LSD) increases mineralocorticoid output by activating a sexually dimorphic transcriptional program

To assess the plasticity of the adrenal cortex in response to environmental requirements, we first stimulated the ZG by feeding male and female mice with a LSD (<0.03% Na^+^) for 2 weeks, starting from 4 weeks until 6 weeks of age (Figure 2a, time point T_1_). To study the ability of the adrenal cortex to reverse the effects of such stimulation, the mice were subjected to a one-week recovery period in which the mice are returned to a standard diet, referred to as T_2_ (Figure 2a). LSD did not change total body weight or adrenal weight of treated mice compared to controls, with the exception of the left adrenal of LSD-fed females, which showed an increase in absolute weight (Figure S6a-b). No major histological changes were observed following treatment or recovery (Figure S6c). As expected, *Cyp11b2* mRNA expression increased significantly after two weeks of LSD, associated with an increase in plasma aldosterone levels (Figure 2b-c), without modification of *Cyp11b1* expression, but increased corticosterone levels in males (Figure 2d-e). After one week recovery (T_2)_, mice fed with a LSD had comparable *Cyp11b2* expression and plasma aldosterone levels to control (Ctrl T_2_) mice (Figure 2b-c). RNAscope analysis confirmed increased mRNA expression of *Cyp11b2* in the ZG of mice following LSD, with no differences in *Cyp11b1* mRNA expression (Figure S7). Consistent with increased *Cyp11b2* expression and plasma aldosterone levels, immunostaining showed a significant increase in AS^+^ cells in the ZG in both male and female mice following LSD and a decrease in AS^+^ cells after recovery, with AS expression returning to levels comparable to Ctrl T_2_ (Figure 2f). Dab2 immunofluorescence revealed no changes in ZG size in both sexes after LSD or recovery (T_2_) (Figure 2g). Steroid profiling of mice under LSD showed an increase of mineralocorticoids as well as steroid precursors, which returned to control values after one-week recovery in both sexes (Figure 2h). PCA of plasma steroids revealed clear separation between control and LSD-fed mice along the first Dimension (Dim1) (Figure 2i), which explained 51.1% in males and 44.9% in females of the total variance, driven mainly by aldosterone, pregnenolone, progesterone, 18-hydroxycorticosterone levels, with a sexually dimorphic contribution of each single steroid (Figure S8a and b). Cell lineage conversion following LSD was investigated in *Cyp11b2^+/Cre^*-*m*T*m*G mice. Overall quantification of GFP^+^ cells within the cortex showed a small, but not significant increase in both male and female mice (Figure S6d). Quantification following segmentation of the adrenal cortex in successive layers revealed an increased number of mGFP^+^ cells in the external layers in males, corresponding to the ZG and compatible with the increased number of AS expressing cells, while in females a lower number of mGFP^+^ cells were observed in the inner layers compared to Ctrl T_1_ mice (Figure S6e).

**Figure 2.**
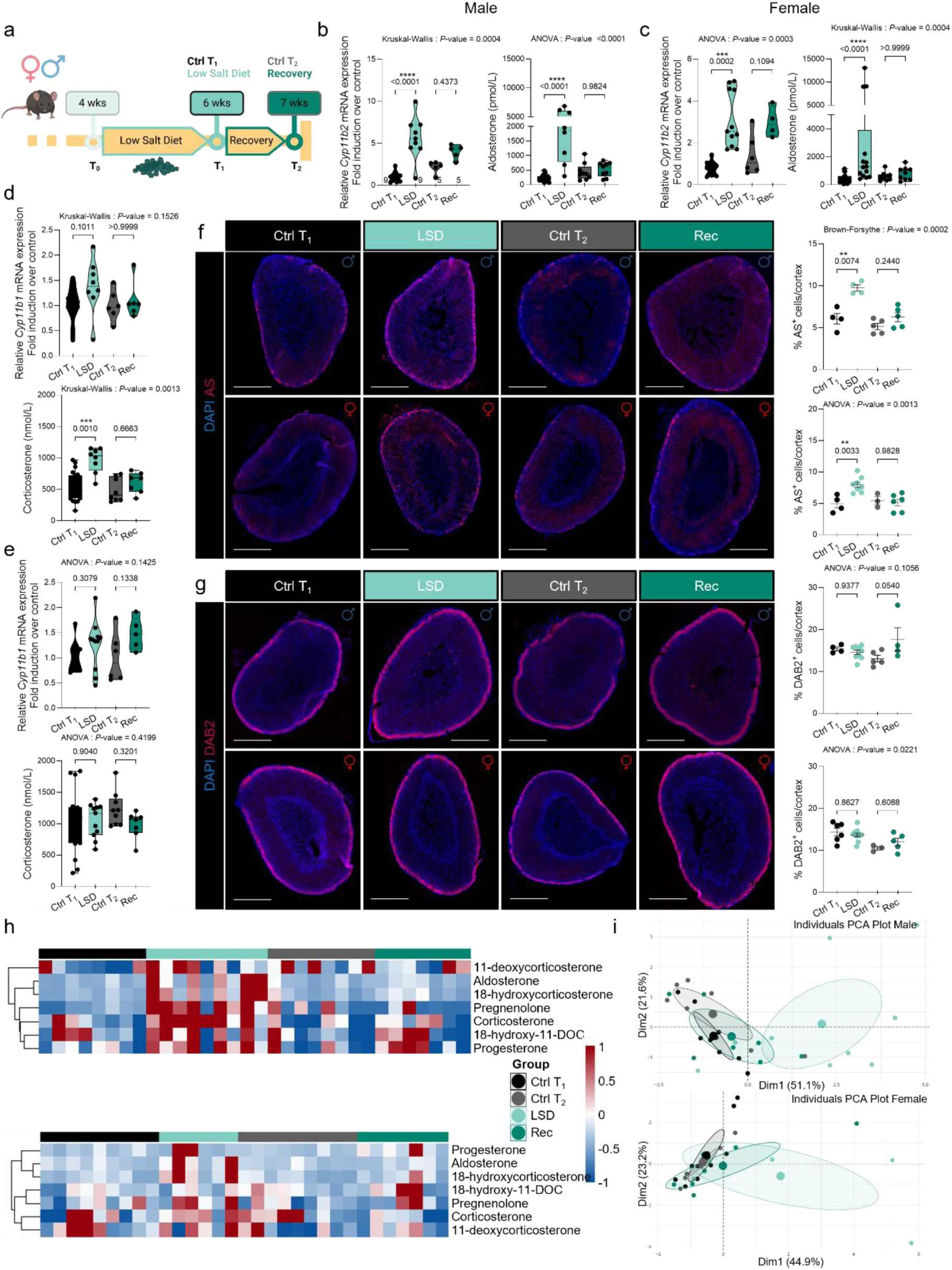
Effect of LSD on the adrenal cortex. **(a).** Schematic representation of the LSD administration regimen from 4 (T_0_) to 6 weeks of age (T_1_), followed by one week recovery to age 7 weeks (T_2_). **(b-c).** Relative *Cyp11b2* mRNA expression measured by RT-qPCR and plasma aldosterone levels measured by LC-MS/MS at T_1_ and T_2_, in male **(b)** and female **(c)** *Cyp11b2^+/Cre^*-*m*T*m*G mice. **(d-e).** RT-qPCR analysis of *Cyp11b1* expression and plasma corticosterone levels measured by LC-MS/MS at T_1_ and T_2_ of **(d)** male and **(e)** female *Cyp11b2^+/Cre^*-*m*T*m*G mice. **(f).** Immunofluorescent labelling of AS (ruby red) and quantification of %AS^+^ cells in the adrenal cortex of male (upper panel) and female (lower panel) mice at T_1_ and T_2_. **(g)**. Immunofluorescent labelling of DAB2 (ruby red) and quantification of the %DAB2^+^ cells in the adrenal cortex of male (upper panel) and female (lower panel) at T_1_ and T_2_. **(h).** Hierarchically clustered heatmaps of adrenal plasma steroid profiles of male (upper pannel) and female (lower pannel) mice at T_1_ and T_2_. **(i).** Principal component analysis (PCA) of adrenal plasma steroid profiles of male (upper pannel) and female (lower pannel) mice at T_1_ and T_2_. P values were obtained with Ordinary one-way ANOVA, followed by Šídák’s multiple comparisons test or Kruskal-Wallis test followed by Dunn’s multiple comparisons test, as indicated. Data are presented as means ± SEM. *P value < 0.05; **P value < 0.01; ***P value < 0.001.

ST data were interrogated to better detect changes in cell components and explore molecular changes underlying the response to LSD. In male mice, LSD induced an increase in clusters corresponding to ZG Adrenal Cortex Progenitors, ZG and the whole ZF (including the ZF, ZF associated macrophages and ZF associated DC clusters) at T_1_ (Figure 3a). In the subsequent recovery week, a ZG Adrenal Cortex Progenitors and ZF clusters were reduced compared to the corresponding control animals, while the other clusters returned to control values (Figure 3c). Analysis of differentially expressed genes (DEG) performed in the ZG and ZF clusters revealed important transcriptional changes in the ZG, with increase in genes involved in steroid hormone biosynthesis and metabolism and ZG cell identity, such as *Cyp11b2, Dab2*, *Hsd3b1*, *Agtr1a*, and mitochondrial function (*Hspd1*, *Echs1*, *Fdx1, Atp5e*) (Figure 3b, f and Table S2). Remarkably, ZF appears to actively contribute to the response to LSD, as revealed by a large number of positive transcriptional changes (395 DEG, 352 upregulated, Figure 3b and Table S2). Among these genes are some that are directly linked with ZF cell identity and function, with increased expression of *Cyp11b1, Akr1b7, Scarb1* and *Fdx1* (Figure 3b and Table S2). Following one week of recovery the transcriptional activity of the ZG reverted to normal, whereas ZF still displays important transcriptional changes compared to Ctrl T_2_ mice (Figure 3d). In female mice, a similar increase of the ZG cluster following LSD was observed, while the ZG Adrenal Cortex Progenitors and ZF clusters were reduced (Figure 3e). Recovery led to re-expansion of ZG Adrenal Cortex Progenitors and decrease of ZG (Figure 3g). LSD was associated with a limited number of transcriptional changes in the female ZG, including increased expression of *Cyp11b2* and *Fdx1*, while *Wnt4* was decreased (Figure 3f and Table S2). In contrast, an important transcriptional repression was observed in the ZF, in particular reduced expression of steroidogenic genes such as *Hsd3b1*, *Akr1b7*, *Scarb1*, and *Cyp21a1* (Figure 3f and Table S2). Recovery was associated to major changes in the ZF but not in the ZG, similar to what observed in males (Figure 3h and Table S2), with a recovery of positive transcriptional regulation (235 upregulated genes) and increased expression of *Hsd3b1*, *Cyp21a1,* and *Cyp11a1*. Overall, a higher number of DEG following LSD were observed in males compared to females, with major differences in the direction of regulation at both T1 and T2 particularly in the ZF (Figure 3i and S9, Table S2). Indeed, gene ontology revealed upregulation of KEGG pathways related to oxidative phosphorylation, steroid biosynthesis, and transcription in the ZF following LSD in males, while in females a majority of pathways was downregulated, including MAPK signalling, oxidative phosphorylation, extracellular matrix (ECM) receptor interaction and neurotrophin signaling (Figure S9 and Table S3). Following recovery, in males the main pathways were downregulated, including oxidative phosphorylation and ECM receptor interaction, while in females recovery was associated with upregulation of ubiquitin mediated proteolysis, mitochondrial function, and apoptosis, and downregulation of calcium signalling, cell adhesion and hedgehog signalling. No modifications of Wnt/β-catenin genes were observed (Figure S6 f-g).

**Figure 3.**
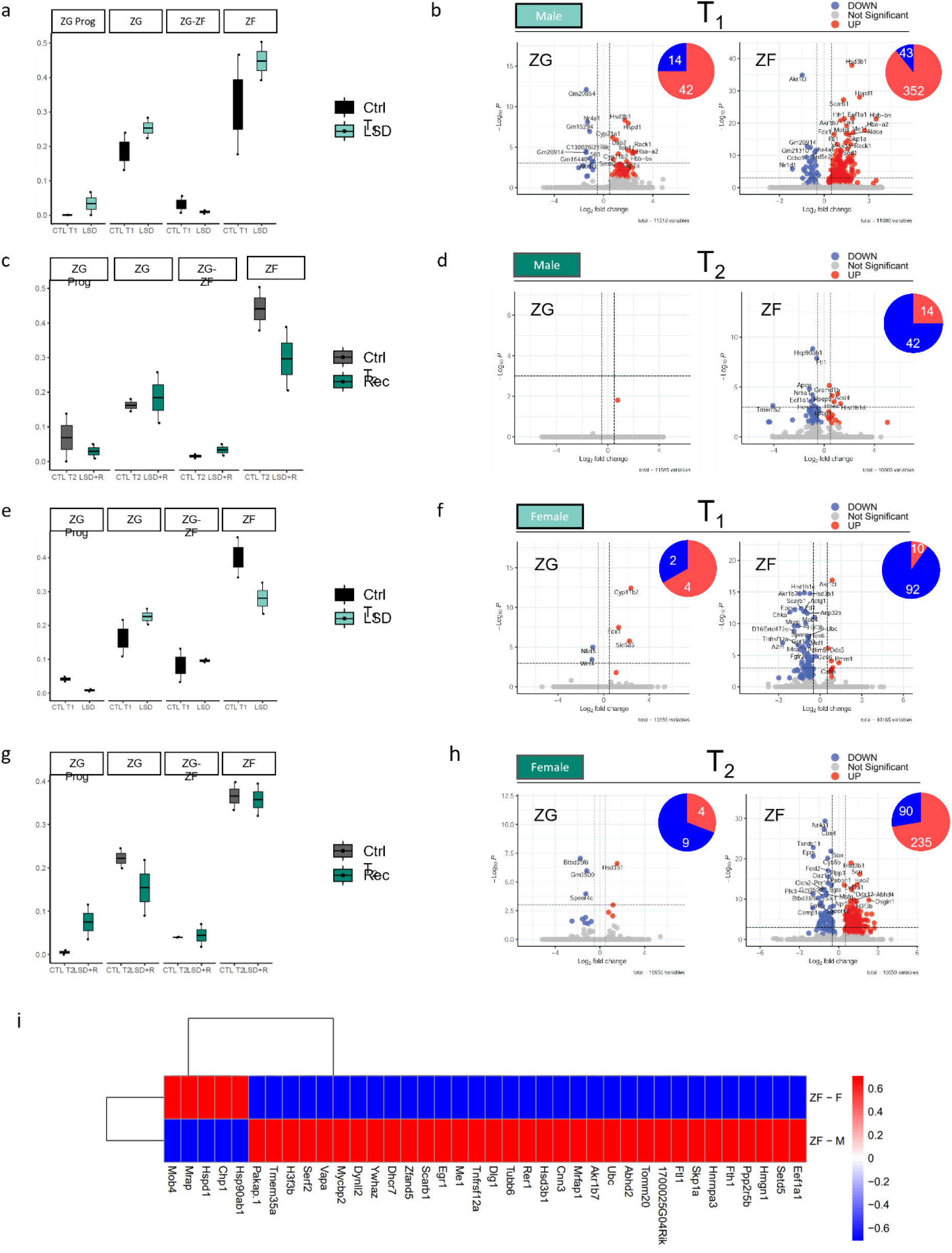
Effect of LSD on zone-specific cell composition and gene expression. **(a,c)**. Cell cluster proportions of male *Cyp11b2^+/Cre^*-*m*T*m*G mice at T_1_ **(a)** and T_2_ **(c)** following a LSD. **(b, d).** Volcano plots of differentially expressed genes (DEGs) in the ZG and ZF cell clusters of male *Cyp11b2^+/Cre^*-*m*T*m*G mice at T_1_ **(b)** and T_2_ **(d)**, with chart pies summarizing the number of DEGs. **(e, g).** Cell cluster proportions of female *Cyp11b2^+/Cre^*-*m*T*m*G mice at T_1_ **(e)** and T_2_ **(g)**. **(f, h)**. Volcano plots of DEGs in the ZG and ZF cell clusters of female *Cyp11b2^+/Cre^*-*m*T*m*G mice at T_1_ **(f)** and T_2_ **(h)**, with chart pie summarizing the DEGs. **(i).** Z-score hierarchical clustering heat map visualization representing the DEGs with opposite regulation in male and female *Cyp11b2^+/Cre^*-*m*T*m*G male mice at T_1_ in the ZF cluster. Volcano plots were generated starting from the unfiltered list of DEGs; upregulated, downregulated and not significant genes are respectively highlighted in blue, red and grey. Only the most significant genes (-Log10P > 1e-03) are labelled.

Altogether, these results suggest that the response of the adrenal cortex to a 2-weeks LSD, starting at 4-weeks of age, leads to increased mineralocorticoid output mediated by an increase in the number of AS^+^ cells and *Cyp11b2* expression in both sexes, with only little increase of ZG cells revealed by ST. Adaptation to a LSD and the following recovery involves a major sexually dimorphic transcriptional response in the ZF, as well as different contributions of cell components of the adrenal cortex.

### Adaptation of the adrenal cortex to a high salt diet (HSD) involves decrease of AS expression in the ZG and transcriptional changes in the ZF

Next, we investigated the effects of an inhibitory stimulus to the ZG on adrenal cortex plasticity, by feeding 4-weeks-old mice a HSD (3.2% Na^+^) for a two-week period, followed by one week recovery (Figure 4a). Treatment induced a significant suppression of aldosterone levels in both sexes (Figure 4b-c), which was accompanied by a significant decrease of *Cyp11b2* mRNA expression in females (Fig. 4c). In males, although the reduction of *Cyp11b2* mRNA expression did not reach statistical significance when measured by RT-qPCR of the entire adrenal cortex (Figure 4b), ST revealed a significant reduction of *Cyp11b2* mRNA expression in the ZG (Table S4). One-week recovery restored *Cyp11b2* mRNA expression and plasma aldosterone levels in both sexes (Figure 4b-c). HSD and recovery did not modify body weight in male mice compared with controls (Figure S10a), while in females there was a significant increase in body weight after HSD that normalized after recovery (Figure S10b). Right and left adrenal weights showed no changes in both males and females at any time point of the diet (Figure S10a-b). H&E staining did not reveal any histological changes at T_1_ nor at T_2_ (Figure S10c). AS immunofluorescent showed a significant reduction of AS^+^ cells in HSD fed mice versus controls (T_1_, Figure 4d), confirmed also by RNAscope targeting *Cyp11b2* (Figure S7), which returned to normal levels after recovery (Figure 4d). These changes were not associated with any modification of ZG thickness, as quantified by the number of Dab2^+^ cells (Figure 4e). HSD also led to increased plasma corticosterone levels (Figure 4f-g), which was accompanied by increased *Cyp11b1* mRNA expression in males only (Figure 4e), as also confirmed by ST (Table S4). These differences disappeared after recovery (Figure 4f-g). Steroid profiling showed an overall decrease in aldosterone and 18-hydroxycorticosterone levels and an increase in steroid precursors following HSD associated with an increase in glucocorticoid biosynthesis (Figure 4h). Recovery led to normalization of the steroid profile of HSD-fed mice. PCA showed clear separation between control and HSD-fed mice along Dim1 (Figure 4i), which explained 57% in males and 55.3% in females of the total variance, driven mainly by 18-hydroxy-11-DOC, pregnenolone, 11-deoxycorticosterone (only in females), corticosterone and progesterone (Figure 4i, S11a-b). Recovery was associated with a slight shift along the Dim2 driven by 18-hydrocorticosterone and aldosterone (Figure 4i and S11a-b). Analysis of *Cyp11b2^+/Cre^*-*m*T*m*G mice revealed an increase of mGFP^+^ cells following LSD in males (Figure S10d), which was distributed over all layers of the adrenal cortex (Figure S10f), suggesting that HSD may lead to increased lineage conversion from ZG to ZF, due to lack of stimulation to the ZG.

**Figure 4.**
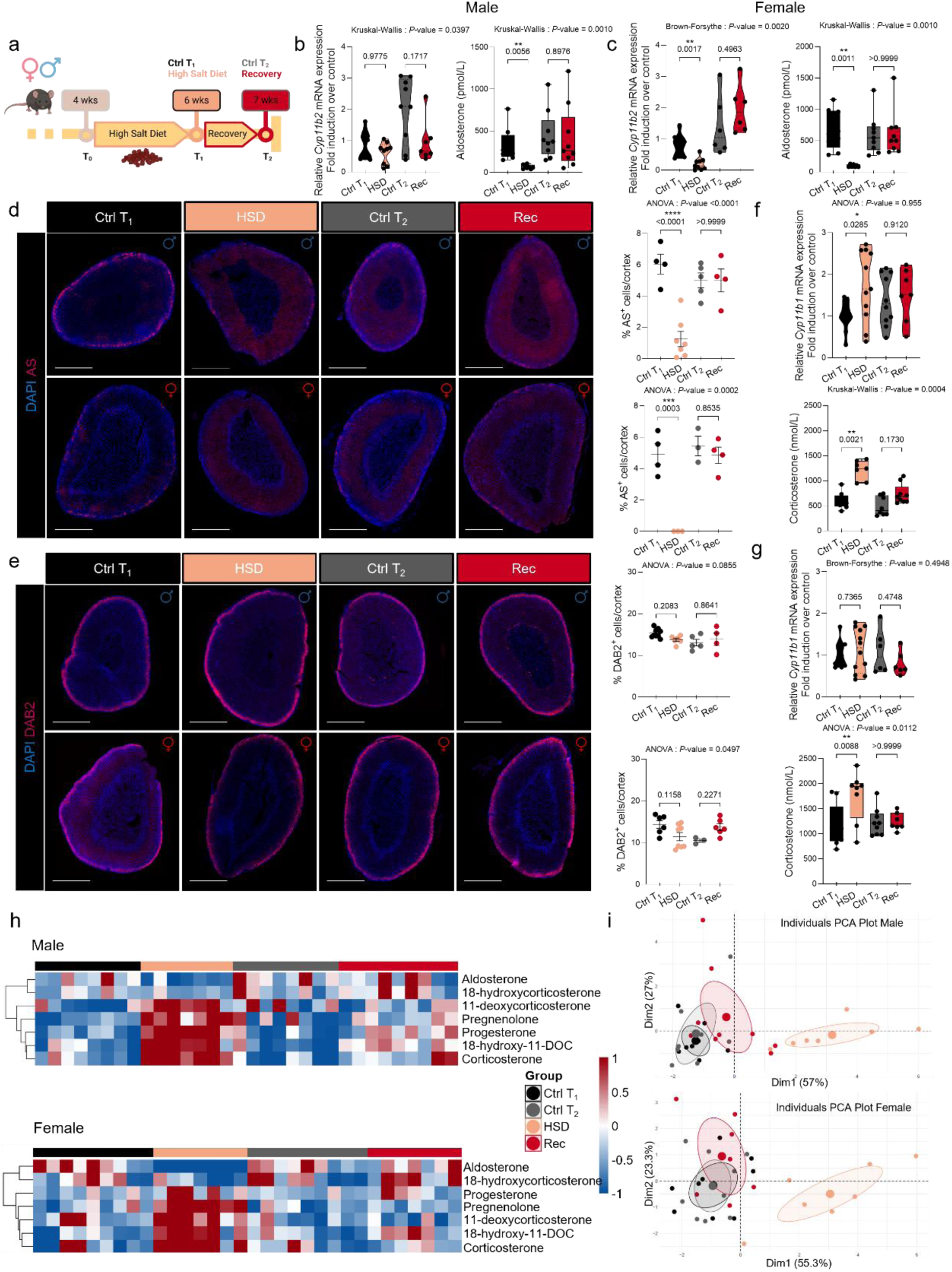
Effect of HSD on the adrenal cortex. **(a)** Schematic representation of the HSD administration regimen from 4 (T_0_) to 6 weeks of age (T_1_), followed by one week recovery to age 7 weeks (T_2_). **(b-c)**. Relative *Cyp11b2* mRNA expression measured by RT-qPCR and plasma aldosterone levels measured by LC-MS/MS at T_1_ and T_2_, in male **(b)** and female **(c)** *Cyp11b2^+/Cre^*-*m*T*m*G mice. **(d)** Immunofluorescent labelling of AS (ruby red) and quantification of the %AS^+^ cells in the adrenal cortex of male (upper panel) and female (lower panel) mice at T_1_ and T_2_. **(e)**. Immunofluorescent labelling of DAB2 (ruby red) and quantification of the %DAB2^+^ cells in the adrenal cortex of male (upper panel) and female (lower panel) at T_1_ and T_2_. **(f-g)**. RT-qPCR analysis for the *Cyp11b1* gene and plasma corticosterone levels measured by LC-MS/MS at T_1_ and T_2_ in male **(f)** and female **(g)** *Cyp11b2^+/Cre^*-*m*T*m*G mice. **(h)**. Hierarchically clustered heatmaps of adrenal plasma steroid profiles of male (upper pannel) and female (lower pannel) mice at T_1_ and T_2_. **(i).** Principal component analysis (PCAs) of adrenal plasma steroid profiles of male (upper pannel) and female (lower panel) mice at T_1_ and T_2_. P values were obtained with Ordinary one-way ANOVA, followed by Šídák’s multiple comparisons test or Kruskal-Wallis test followed by adjusted with Dunn’s multiple comparisons test or Brown-Forsythe ANOVA test followed by Dunnett’s T3 multiple comparisons test, as indicated. Data are presented as means ± SEM. *P value < 0.05; **P value < 0.01; ***P value < 0.001.

ST showed only a slight reduction in the ZG component, but revealed an enrichment in the ZG Adrenal Cortex Progenitors and ZF clusters compared with Ctrl T_1_ in both sexes (Figure 5a, e). After 1 week recovery, a slightly reduced ZG Adrenal Cortex Progenitor cell cluster was observed in males compared to control T_2_, with normalization of the other cell clusters (Figure 5c). In females, recovery was followed by expansion of the ZG-ZF cluster and a minor expansion of ZG and ZF compared with Ctrl T_2_ mice (Figure 5g). DEG analysis showed only a limited number of transcriptional changes after HSD or recovery in the ZG (Figure 5b, d, f, h and Table S4), including downregulation of *Cyp11b2* and *Kcnk9* (which did not reach statistical significance after Bonferroni correction in females). Remarkably, ZF once again showed significant gene expression changes after HSD (Figure 5b, f), as well as after recovery (Figure 5d, h), with important sex differences in the genes and direction of regulation (Figure S12 and Table S4). In males, upregulated genes following HSD are involved in steroid metabolism and mitochondrial function such as *Hsd3b1*, *Fdx1*, *Cyp11a1*, *Cyp11b1*, *Me1*, *Hspd1*, *Tspo*, *Sod1*, as well as ion homeostasis (*Kcnn2*, *Cacna2d1*, *Cacna1c*). In females, upregulated genes are also involved in steroidogenesis and mitochondrial metabolism (*Fdx1*, *Creb3*, *Scarb1*). Similar to what observed with LSD, only a limited number of genes were commonly regulated in males and females (Figure S12). GSEA again revealed opposite regulations in males and females, with KEGG terms related to oxidative phosphorylation being upregulated and neuroactive ligand receptor being downregulated following HSD in males, while females showed upregulation of neuroactive ligand receptor and downregulation of steroid biosynthesis; retinol metabolism was downregulated in both sexes (Table S2). After recovery, oxidative phosphorylation was the most significantly downregulated term in males, with upregulation of steroid hormone biosynthesis, while females showed mostly downregulation of KEGG terms, including calcium signalling, metabolism by cytochrome p450 and cell adhesion molecules. HSD induced only minor modifications of genes of the Wnt/β-catenin pathway (Figure S10e-f).

**Figure 5.**
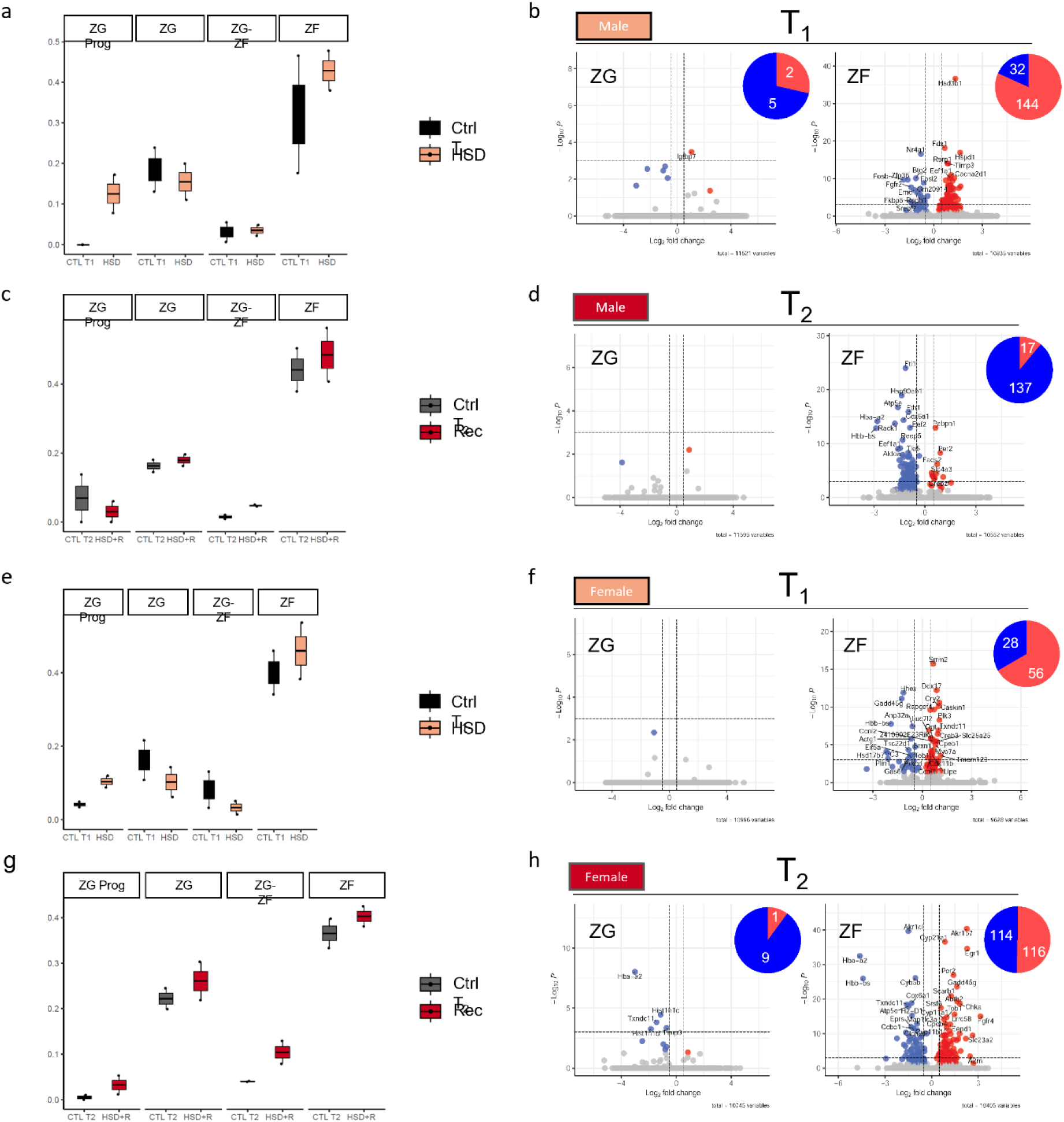
Effect of HSD on zone-specific gene expression and cell composition. **(a,c)**. Cell cluster proportions of male *Cyp11b2^+/Cre^*-*m*T*m*G mice at T_1_ **(a)** and T_2_ **(c)**. **(b, d).** Volcano plots of differentially expressed genes (DEGs) in the ZG and ZF cell clusters of male *Cyp11b2^+/Cre^*-*m*T*m*G mice at T_1_ **(b)** and T_2_ **(d)**, with chart pies summarizing the number of DEGs. **(e, g).** Cell cluster proportions of female *Cyp11b2^+/Cre^*-*m*T*m*G mice at T_1_ **(e)** and T_2_ **(g)**. **(f, h)**. Volcano plots of DEGs in the ZG and ZF cell clusters of female *Cyp11b2^+/Cre^*-*m*T*m*G mice at T_1_ **(f)** and T_2_ **(h)**, with chart pie summarizing the DEGs. Volcano plots were generated starting from the unfiltered list of DEGs; upregulated, downregulated and not significant genes are respectively highlighted in blue, red and grey. Only the most significant genes (-Log10P > 1e-03) are labelled.

Altogether these data suggest that physiological response of aldosterone following a 2-week HSD, starting at 4 weeks of age, is mediated by a decrease in the number of AS expressing cells, reduced *Cyp11b2* expression, but only minor changes in ZG size. The response is associated with a sexually dimorphic transcriptional program involving mainly the ZF. In males, a small increase in GFP^+^ cells distributed over all cortex layers was observed, suggesting that HSD may lead to accelerated lineage conversion from ZG to ZF.

### Dexamethasone (DEX) treatment affects ZG function and accelerates ZG to ZF cell lineage conversion

To investigate whether challenges to the HPA axis similarly affect the interaction between adrenocortical zones, male and female mice underwent two weeks of DEX treatment starting at 4 weeks of age, followed by three weeks of recovery (Figure 6a). The efficacy of the treatment was confirmed by showing a significant reduction in *Cyp11b1* mRNA levels after two weeks of treatment, which returned to control levels after three weeks of recovery (T_2_) (Figure 6b-c). Plasma corticosterone levels were significantly decreased following DEX treatment, returning to normal after recovery in both sexes (Figure 6b-c). DEX treated mice were found to have lower body weight compared with controls, with weight regained in females after recovery, but not in males (Figure S13a-b). This was also associated with a significant reduction in both right and left adrenal weights following DEX treatment, which reversed upon recovery, however without reaching control levels in male mice (Figure S13a-b). H&E staining shows an atrophic ZF with reduced cell density following treatment, with 3-weeks recovery allowing to restore adrenal cortex histology comparable to control (Figure S13c). RNAscope detection of *Cyp11b1* confirmed significant atrophy of the ZF after DEX treatment (Figure S7). When referred to the entire adrenal cortex, an increased number of AS^+^ cells were measured by immunofluorescence in male mice following DEX treatment (Figure 5d), which was associated with a non-significant increase in ZG thickness measured by the number of Dab2^+^ cells (Figure 5e). This increase was however due to a decrease in the total adrenal cortex size due to regression of the ZF following DEX treatment, as indicated by quantification of AS^+^ cells in the first 4 layers of the cortex only (Figure S13d). In females, no major changes in ZG size or number of AS^+^ cells were observed, nor was any difference found after the recovery phase (Figure 5d-e).

**Figure 6.**
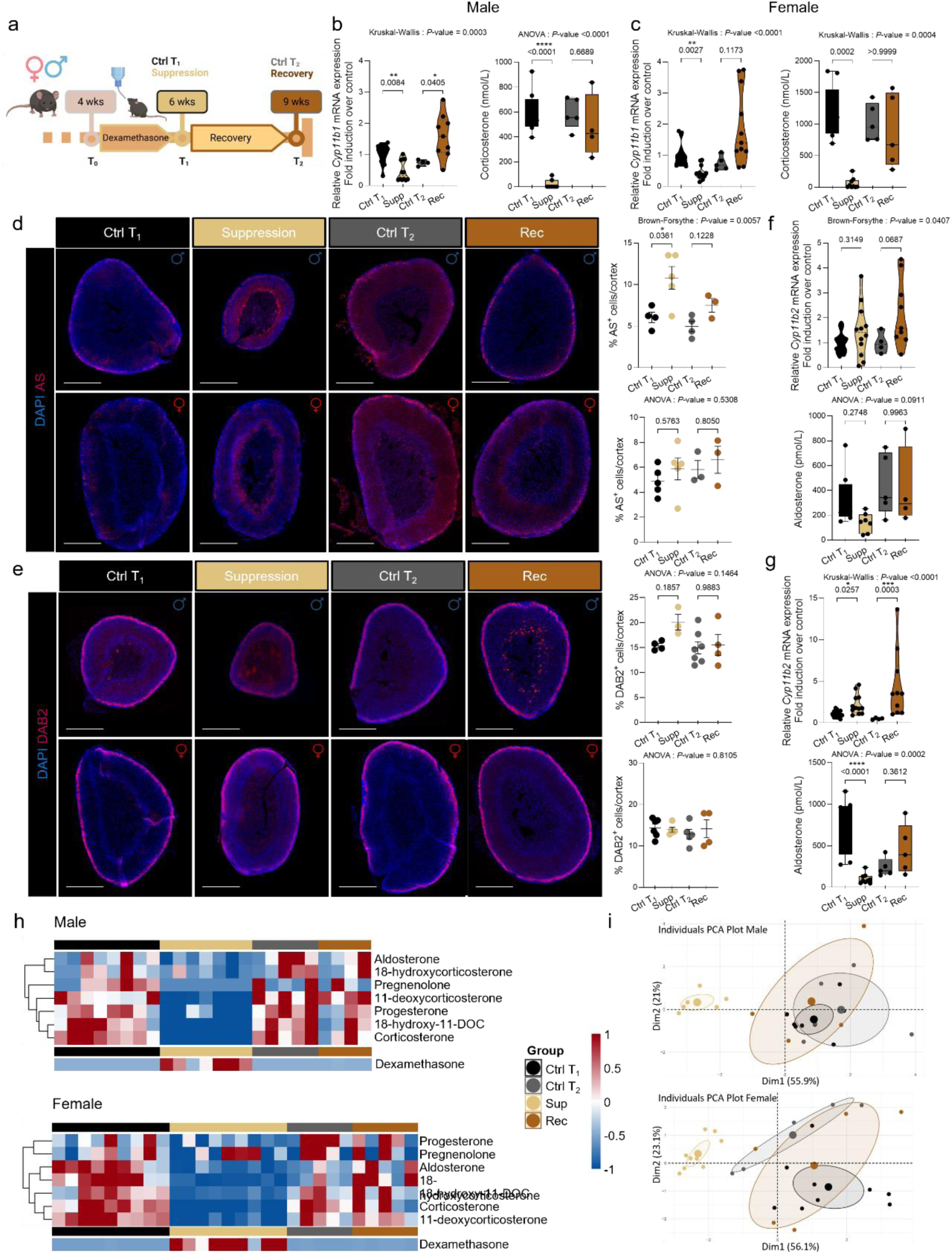
Effect of dexamethasone treatment on the adrenal cortex. **(a)** Schematic representation of the dexamethasone administration regimen from 4 (T_0_) to 6 weeks of age (T_1_), followed by 3 weeks recovery from age 6 weeks to 9 weeks (T_2_). **(b-c)**. Relative *Cyp11b1* mRNA expression measured by RT-qPCR and plasma corticosterone levels measured by LC-MS/MS at T_1_ and T_2_, in male **(b)** and female **(c)** *Cyp11b2^+/Cre^*-*m*T*m*G mice. **(d)** Immunofluorescent labelling of AS (ruby red) and quantification of the %AS^+^ cells in the adrenal cortex of male (upper panel) and female (lower panel) at T_1_ and T_2_. **(e)**. Immunofluorescent labelling of DAB2 (ruby red) and quantification of the %DAB2^+^ cells in the adrenal cortex of male (upper panel) and female (lower panel) at T_1_ and T_2_. **(f-g)**. RT-qPCR analysis for the *Cyp11b2* gene and plasma aldosterone levels measured by LC-MS/MS at T_1_ and T_2_ in male **(f)** and female **(g)** *Cyp11b2^+/Cre^*-*m*T*m*G mice. **(h)**. Hierarchically clustered heatmaps of adrenal plasma steroid profiles of male (upper panel) and female (lower panel) mice at T_1_ and T_2_. Measures of plasma dexamethasone concentrations are indicated below for each animal. **(i).** Principal component analysis (PCA) of adrenal plasma steroid profiles of male (upper panel) and female (lower panel) mice at T_1_ and T_2_. P values were obtained with Ordinary one-way ANOVA, followed by Šídák’s multiple comparisons test or Kruskal-Wallis test followed by adjusted with Dunn’s multiple comparisons test or Brown-Forsythe ANOVA test followed by Dunnett’s T3 multiple comparisons test, as indicated. Data are presented as means +/-SEM. *P value < 0.05; **P value < 0.01; ***P value < 0.001.

*Cyp11b2* expression was not different between treated animals and controls in males, but slightly increased in females following suppression and after recovery. In females, plasma aldosterone levels following DEX treatment were significantly reduced, but normalised after the 3-weeks recovery (Figure 6f-g), suggesting a compensatory increase of *Cyp11b2* mRNA expression to compensate for reduced aldosterone production following DEX treatment. In male mice, although a reduction of plasma aldosterone was observed, this did not reach statistical significance. Steroid profiling revealed an overall decreased production of glucocorticoids and mineralocorticoids as well as steroid precursors following DEX treatment in both sexes, which returned to normal values after the recovery weeks (Figure 6h). PCA of plasma steroids revealed clear separation between controls and DEX-treated mice along the first Dimension (Dim1), which explained 55.9% in males and 56.1% in females of the total variance, driven mainly by a reduction in corticosterone, 11-deoxycorticosterone, 18-hydroxy-11-DOC, progesterone (only in males) and 18-hydrocorticosterone (only in females) (Figure 6i, S14a-b). Steroid profiles completely normalised following 3 weeks of recovery (Figure 6i). Analysis of cell lineage conversion in *Cyp11b2^+/Cre^-mTmG* mice showed a slight increase in total number of GFP^+^ cells in the adrenal cortex in males, which was evenly distributed over all quantified layers (Figure S13e).

ST revealed a major transcriptional induction in the ZG in male mice (Figure 7a, Table S5), with 104 upregulated genes, together with an increase of the ZG Adrenal Cortex Progenitors and the ZG cluster, and a completely suppressed ZF cluster as expected. DEG analysis did not reveal any regulated genes in the ZF due to the severe atrophy following treatment. Among the most upregulated genes in the ZG were *Hsd3b1*, as well as *Fkbp4*, *Mc2r*, coding for the ACTH receptor, *Wnt5a*, a component of the Wnt/β-catenin pathway, and *Dlk*. *Nr4a1*, coding for NGFI-B, was one of the 5 downregulated genes, together with *Akr1b7* and *Vegfa*. Upregulated pathways included lysosome activity, PPAR signalling, and ABC transporters, which are associated with enhanced cellular metabolism and signalling, while the downregulated functions, including unsaturated fatty acid biosynthesis, steroid biosynthesis, retinol metabolism, indicate suppression of steroid metabolism (Figure S15). In female mice, DEX suppressed the ZF cluster as well as the ZG cluster (Figure 7e). DEG analysis following DEX treatment in females did show only slight changes in the ZG, but significant transcriptional regulation in the ZF (Figure 7d). Among the most upregulated genes were *Apoe*, *Nrep* and *Prkcd*, while down-regulated genes included *Cdkn1c*, *Kcnk3*, *Npr1*, *Cyp21a1*, and *Nr4a2*. In contrast to males, pathways related to steroid biosynthesis were upregulated in both ZG and ZF (Figure S15). RT-qPCR showed also activation of components of the Wnt/β-catenin pathways after DEX, which have suggestive significance in ST, such as *Wnt4* and *Lef1* in males, whereas increased expression of *Axin2* in both sexes was detected only by RT-qPCR (Figure 7c, f). In male mice, expression of *Tcf3* and *Axin2* were increased after recovery compared to Ctrl T2 (Figure 7c). DEX also induced an important increase of the Macrophage cluster mainly in male mice (Figure 7b). Those macrophages were localised at the border between the ZF and the medulla (Figure S4), as shown by CD68 immunostaining of adrenals from Cyp11b2^+/Cre^-mTmG mice (Figure 7g). Those cells were lipid-laden, as shown by Oil Red O staining (Figure 7h), suggesting a “foam cell” phenotype. Finally, immunofluorescent staining of Ki67, a marker of proliferation, showed a strong decrease of Ki67^+^ nuclei in the adrenal cortex after DEX in both sexes, the only cells in active proliferation being located at the interface between the ZG and ZF (Figure 7i).

**Figure 7.**
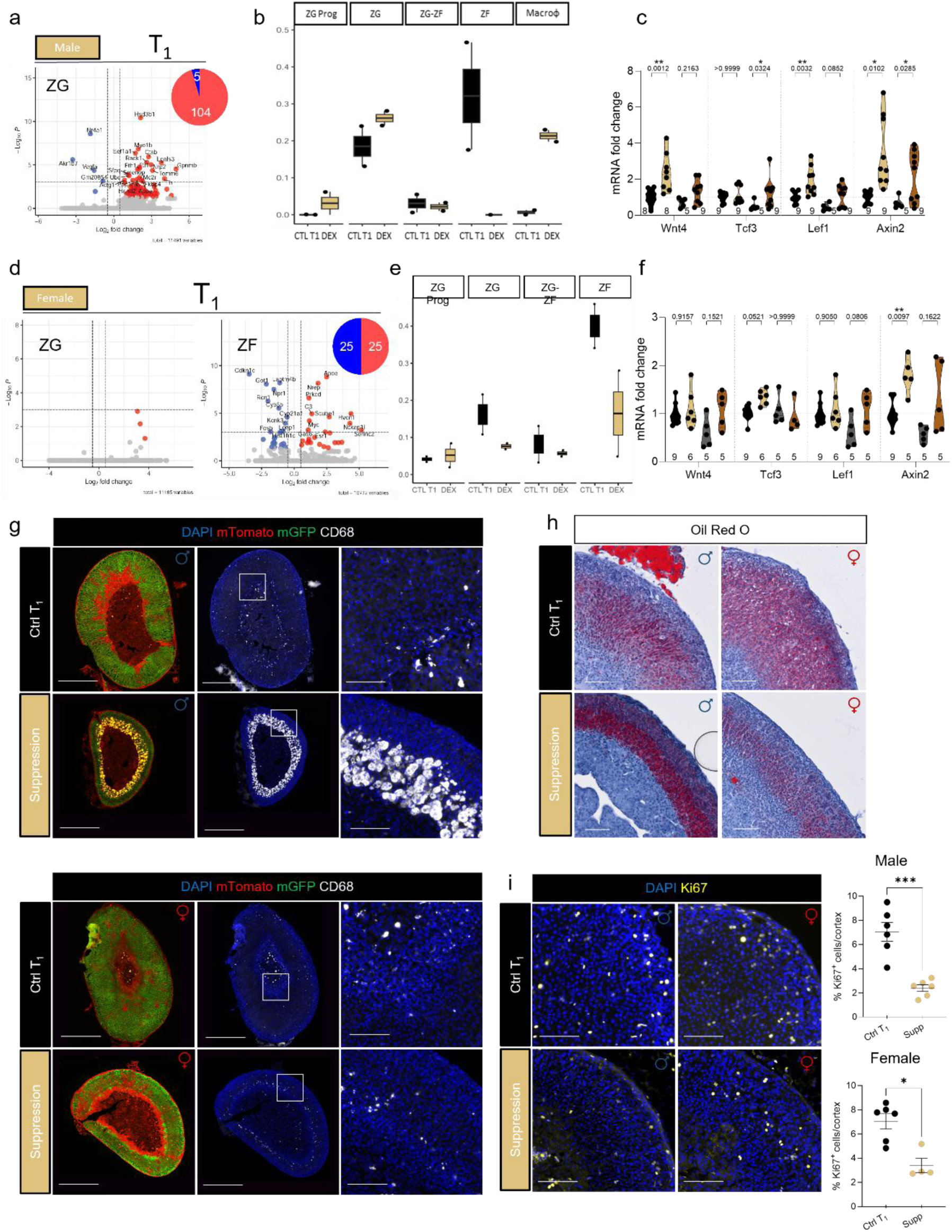
Effect of DEX treatment on zone-specific gene expression and cell composition. **(a,d)**. Volcano plot of DEGs in the ZG cell cluster of male **(a)** and in the ZG and ZF cell clusters of female **(d)** *Cyp11b2^+/Cre^*-*m*T*m*G mice at T_1_, with chart pies summarizing the number of DEGs. **(b,e).** Cell cluster proportions of *Cyp11b2^+/Cre^*-*m*T*m*G male **(b)** and female **(e)** mice at T_1_. **(c, f)** Wnt4, Tcf3, Lef1 and Axin2 mRNA expression on male **(c)** and female **(f)** mice at T_1_ and T_2_. **(g).** Coimmunofluorescence of mGFP (green) with mTomato (red) and CD68 (white) of *Cyp11b2^+/Cre^*-*m*T*m*G male (upper panel) and female (lower panel) mouse adrenals treated with dexamethasone (suppression) vs controls at T_1_. Scale Bar: 500 μm left panels with entire adrenals and 100 μm left pannel. White squares represent the magnification area in the lower panel. **(h).** Oil Red O staining of *Cyp11b2^+/Cre^*-*m*T*m*G male (left panel) and female (right panel) mouse adrenals treated with dexamethasone (suppression) vs controls at T_1_. Scale Bar: 100 μm. **(i).** Immunofluorescent labelling of Ki67 (yellow) and quantification of the %Ki67^+^ nuclei in the adrenal cortex of *Cyp11b2^+/Cre^*-*m*T*m*G (upper panel) male and (lower panel) female mouse adrenals treated with dexamethasone vs controls at T_1_, and quantification of the %Ki67^+^ nuclei in the adrenal cortex. Scale Bar: 100 μm. P values were obtained with Ordinary one-way ANOVA, followed by Šídák’s multiple comparisons test or Kruskal-Wallis test followed by adjusted with Dunn’s multiple comparisons test between the different time points. Data are shown as mean ± SEM. *P value < 0.05; **P value < 0.01; ***P value < 0.001.

Altogether these results suggest that DEX treatment, similar to salt diets, affects both ZG and ZF, with an overall suppression of both glucocorticoid and mineralocorticoid axes as well as sexually dimorphic changes in gene expression, zone expansion and transdifferentiation.

### LSD, HSD and DEX affect ZX regression and lineage conversion in female mice

Since our results indicated a certain degree of lineage conversion between the ZF and the ZX in female mice, we investigated whether the ZX is affected by a LSD, HSD or DEX treatment in female mice. The number of 20αHSD^+^ cells did not change significantly following a HSD or LSD (Figure 8a). With the resolution of ST, an increase in the ZX cluster was observed following LSD compared to controls (Figure 8b). This was associated with a lower number of GFP^+^/20αHSD^+^ cells in LSD mice compared with controls (Figure 8a and c). No major changes were observed following a HSD. While the number of 20αHSD^+^ cells decreases between 6 (Ctrl T_1_, Figure 8a) and 7 weeks (Ctrl T_2_, Figure 8d and 8g) of age, no such decrease is observed in mice following a LSD after 1 week of recovery (Figure 8g) and the number of double positive GFP^+^/20αHSD^+^ cells appears slightly, although not significantly, increased. Similar changes were also observed following a HSD (Figure 8g). These changes are confirmed by ST, with a larger ZX component at T2 following LSD or HSD (Figure 8f).

**Figure 8.**
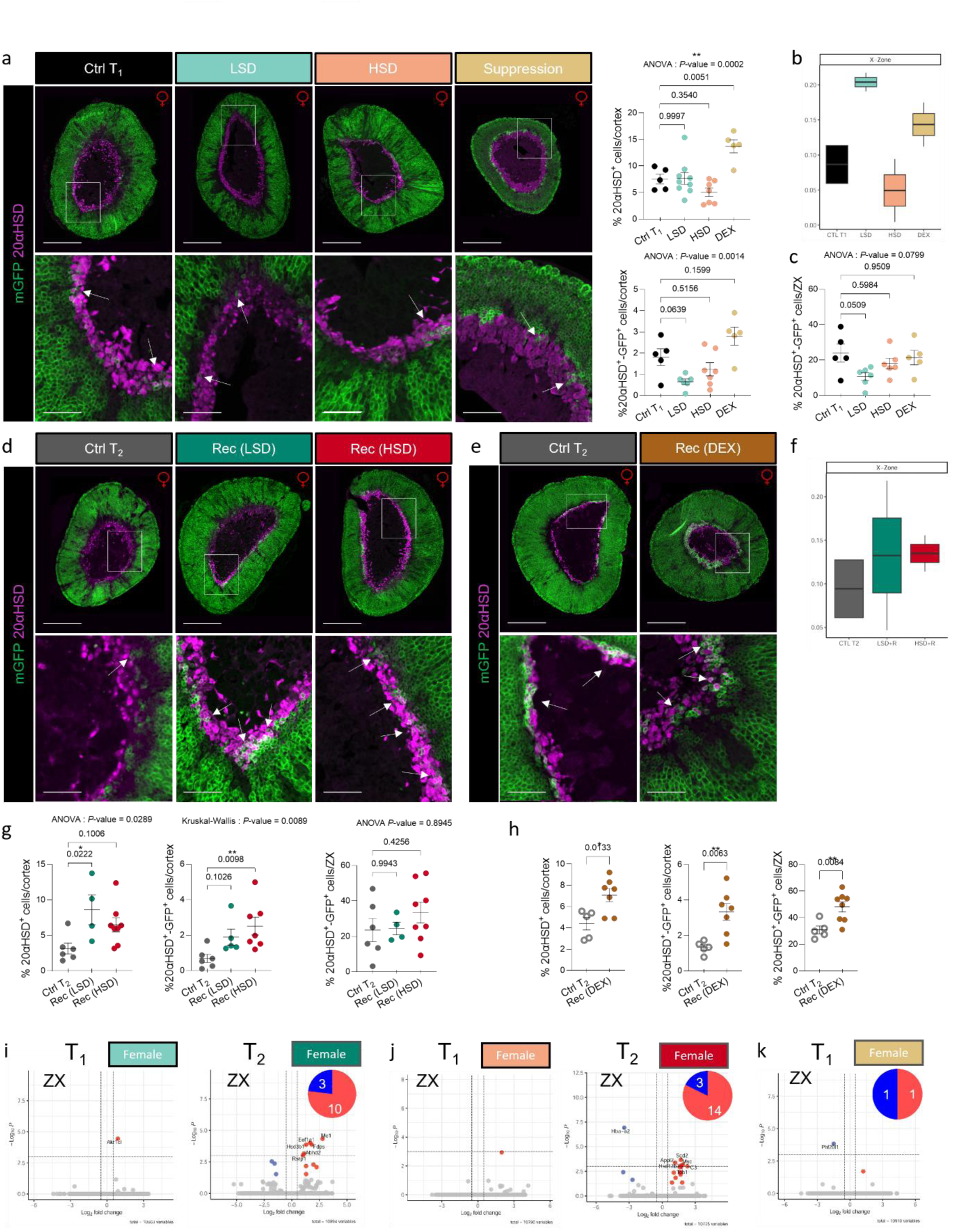
X-Zone plasticity in response to pathophysiological challenges. **(a)**. Coimmunofluorescence of mGFP (green) and 20αHSD (purple) of *Cyp11b2^+/Cre^*-*m*T*m*G female mice that underwent LSD, HSD and DEX treatment vs Ctrl T_1_ and quantification of 20αHSD^+^and mGFP and 20αHSD double-positive cells in the adrenal cortex. mGFP and 20αHSD double-positive cells are indicated by white arrows in the lower panel magnified image. Scale Bar: 500 μm upper panels with entire adrenals and 100 μm lower panels. White squares represent the magnification area in the lower panel. **(b)**. ST ZX cluster proportions of *Cyp11b2^+/Cre^*-female mice that underwent LSD, HSD and DEX treatment vs Ctrl T_1_. **(c).** Quantification of the mGFP and 20αHSD double-positive cells in the X-zone of *Cyp11b2^+/Cre^*-*m*T*m*G female mice that underwent LSD, HSD and DXM vs Ctrl T_1_. **(d)**. Coimmunofluorescence of mGFP (green) and 20αHSD (purple) of *Cyp11b2^+/Cre^*-*m*T*m*G female mice that underwent LSD, HSD after a 1-week recovery period vs Ctrl T_2_ (7 weeks old) mice. mGFP and 20αHSD double-positive cells are denoted by white arrows in the lower panel magnified image. The Ctrl T_2_ image has been used on Figure 1d. Scale Bar: 500 μm upper panels with entire adrenals and 100 μm lower panel. White squares represent the magnification area in the lower panel. **(e).** Coimmunofluorescence of mGFP (green) and 20αHSD (purple) of *Cyp11b2^+/Cre^*-*m*T*m*G female mice treated with DEX followed by 3-weeks recovery vs Ctrl T_2_ (9 weeks old) mice. mGFP and 20αHSD double-positive cells are denoted by white arrows in the lower panel magnified image. Scale Bar: 500 μm upper panels with entire adrenals and 100 μm lower panel. White squares represent the magnification area in the lower panel. **(f).** ST ZX cluster proportions of *Cyp11b2^+/Cre^*-female mice that underwent LSD, HSD and 1-week recovery period vs Ctrl T_2_ (7 weeks old) mice. **(g)**. Quantification of 20αHSD^+^, mGFP and 20αHSD double-positive cells in the adrenal cortex and X-zone of *Cyp11b2^+/Cre^*-*m*T*m*G female mice that underwent LSD, HSD and 1-week recovery period vs Ctrl T_2_ (7 weeks old) mice. **(h)**. Quantification of 20αHSD^+^, mGFP and 20αHSD double-positive cells in the adrenal cortex and in the X-zone of *Cyp11b2^+/Cre^*-*m*T*m*G female mice treated with DEX followed by 3-weeks recovery vs Ctrl T_2_ (9 weeks old) mice. P values were obtained with Unpaired t test. **(i,j,k).** Volcano plot of DEGs in the X-zone cell cluster of *Cyp11b2^+/Cre^*-*m*T*m*G female mice of mice fed a LSD **(i)** and HSD **(j)** at T_1_ and T_2,_ and following DEX cohort at T_1_ **(k)**, with chart pies summarizing the number of DEGs. P values were obtained with Ordinary one-way ANOVA, followed by Šídák’s multiple comparisons test or Kruskal-Wallis test followed by adjusted with Dunn’s multiple comparisons test or Brown-Forsythe ANOVA test followed by Dunnett’s T3 multiple comparisons test, as indicated. Data are presented as means +/-SEM. *P value < 0.05; **P value < 0.01; ***P value < 0.001.

In mice treated with DEX, the ratio of the ZX 20αHSD^+^ cells over the adrenal cortex is increased (Figure 8a), as also confirmed by ST (Figure 8b). Although DEX treated mice reduce the size of their ZX at 9 weeks of age after 3-weeks recovery, the number of 20αHSD^+^/ZX cells is significantly higher in treated mice than in controls at T_2_ (Figure 8h). The lineage conversion process is also significantly altered by DEX treatment, with an increased number of GFP^+^/20αHSD^+^ cells following recovery, representing around 50% of ZX cells (Figure 8h).

Analysis of DEG in the ZX following treatments showed transcriptional changes mainly following recovery after LSD or HSD (Figure 8i, Table S6), with upregulation of genes involved in steroid metabolism *Me1* and *Hsd3b1* and downregulation of *Nr4a1* following LSD, and upregulation of genes involved in cholesterol and lipid metabolism following HSD (*Scd2*, *Hsd17b7*, *Cyp51*, *Msmo1*, *Fdps*).

Altogether, these results indicate that challenges affecting the ZG and ZF, such as salt diets and modulation of the HPA axis, also affect the ZX, in particular its regression with time and cell lineage conversion rate from the ZF.

## Discussion

In this work we have explored how environmental cues affecting mineralocorticoid output, such as salt diets, modulate adrenal cortex homeostasis, cell lineage conversion and zone-specific transcriptional landscape. To this purpose, we have used multiple complementary morphological, functional and molecular investigations on a newly developed *Cyp11b2^cre^* mouse model. This model, in which the Cre recombinase is inserted into the 3’UTR of the *Cyp11b2* gene, conserves normal ZG function, allowing its use to explore mineralocorticoid pathophysiology. We show that *Cyp11b2* expression is detectable as early as P1 in different areas of the ZG and a progressive reduction of ZG size and evolution of cell components of the adrenal cortex with age. Cell lineage conversion in *Cyp11b2^+/Cre^-mTmG* mice showed the typical progressive transdifferentiation of ZG into ZF cells over time, progressing into the ZX in females, revealing a previously unrecognized connection between ZF and ZX cells in adult mice. A 2-weeks HSD or LSD, starting from 4 weeks of age, modulates mineralocorticoid output mainly by regulating steroidogenic gene expression, with little modifications of the number of ZG cells, but induces a major, sexually dimorphic, cellular and transcriptional remodelling in the ZF, which is also the major hallmark of recovery. A similar reciprocal response of the ZG is observed when challenging the ZF by DEX treatment, indicating a functional and molecular interaction between ZG and ZF to adapt to pathophysiological challenges.

Our study has allowed gaining insight into major processes involved in adrenal growth and reveals dynamic changes of adrenal cortex cell composition over time that are associated with functional and structural evolution and hormone production. GFP^+^ cells were present in large areas of the adrenal surface already at P1, reflecting expression of *Cyp11b2,* and expanded into the entire ZG by two weeks of age, with cell lineage conversion proceeding into the entire ZF between 6 and 9 weeks of age in both males and females. These data are different from a previous model, in which the Cre recombinase was inserted into exon 2 of the *Cyp11b2* gene^19^. This may be due to a certain degree of *Cyp11b2* haploinsufficiency in the previous model, which, although without overt phenotype during postnatal life, may have modified *Cyp11b2* expression during embryonic development and subsequent expansion of AS expressing cells. We observed a progressive reduction of Dab2^+^ cells between 2 and 6 weeks of age, which was paralleled by a reduction of plasma aldosterone levels. This is in agreement with the evolution of aldosterone levels in humans, where high aldosterone levels are observed in newborns, progressively decreasing with age^25–27^. Remarkably, our data show that lineage conversion in female mice progresses into the ZX, as evidenced by the coexpression of mGFP and 20αHSD, a ZX marker, in female 6- and 9-weeks old *Cyp11b2^+/Cre^-mTmG* mice. Previous lineage tracing experiments have shown that the mouse adult cortex derives from precursor cells in the foetal cortex, demonstrating a direct link between the foetal and adult zones during development^14^. It has further been postulated that the foetal zone derives from the definitive zone through centripetal migration of cells^28^. Consistent with these models, recent data suggest a shared progenitor between the definitive and foetal zones in humans and cynomolgus monkey^29^. This “adrenal primordium” can differentiate into the definitive zone or the foetal zone through a direct pathway; in addition, an indirect pathway has been identified in which cells from the definitive zone differentiate further into the foetal zone, which thus represents the final stage of lineage conversion^29^. Our data indicate that, unlike the faster transdifferentiation observed between ZG and ZF, final lineage conversions from the ZF to the ZX proceeds much slower and never fully completes the process. Further investigations employing specific cell lineage tracing models may allow to further delineate this process.

Our results also suggest that the response of the ZG to a 2-week salt challenge with a LSD or HSD does not involve major modifications of ZG structure and cell lineage conversion, with only little modifications in ZG size, but is rather mediated by an effect on *Cyp11b2* expression, both by modulating its expression levels as well as the number of AS expressing ZG cells. Only a small increase in the ZG cluster was revealed by the resolution of ST. This is consistent with previous observations by Nishimoto et al, in which a short salt challenge to rats did not expand the ZG^24^, while other models showed increased ZG following a low salt diet^23^. It is interesting to note that we show a progressive reduction of ZG size until 4 weeks of age, suggesting a reduced proliferative capacity of the adrenal ZG afterwards. This supports a hypothesis whereby timing of salt challenge, in addition to duration, may influence the expansive capacity of the ZG. Remarkably, we reveal a sexually dimorphic cellular and transcriptional adaptation to salt challenges, as well as interactions between the different cell layers of the adrenal cortex. Indeed, the resolution of ST allows to appreciate a small increase or decrease of the ZG cluster following a LSD or HSD respectively, associated with changes in the expression genes related to ZG cell identity and function. In addition, major modifications are observed in the ZF, which responds to salt challenges with changes in size and a large transcriptional and sexually dimorphic remodelling, which also contributes to recovery of the adrenal cortex after returning to normal salt diet. RAS modulation did not appear to affect the Wnt/β-catenin pathway in our experimental conditions. Previous studies have shown that changes in Wnt/β-catenin impact *Cyp11b2* expression and aldosterone production, as described in the Wnt4-deficient model (Wnt4^-/-^)^30^ and in studies using constitutive β-catenin activation (β-cat-GOF model)^31,32^. Our study indicates that, conversely, signals that increase (LSD) or decrease (HSD) *Cyp11b2* expression and aldosterone production do not affect Wnt/β-catenin signalling. Thus, salt diets do not appear to alter the main signalling pathway involved in maintaining the cellular identity and differentiation of the ZG, suggesting that RAS and Wnt/β-catenin pathways may regulate ZG through different, complementary mechanisms and that AS itself may be a component of ZG identity, whose expression is not only regulated towards maintaining salt and water homeostasis but may also play a role in more complex cellular functions.

DEX suppression is well known to suppress the endogenous HPA axis, inducing major changes in the adrenal cortex, including ZF atrophy with increased apoptosis, decreased *Cyp11b1* expression and corticosterone production. Recovery following DEX suppression involves cortical sonic hedgehog (Shh)^+^ progenitor cells and Shh-responsive capsular cells, as well as WNT signalling^22^. Here we show that DEX treatment induces major ZF atrophy and infiltration of macrophages, which is more pronounced in males, as well as a general suppression of corticosteroid production, with reduced aldosterone levels particularly evident in females, which may be related to the role of ACTH in regulating aldosterone production. However, beyond this endocrine regulation, DEX suppression also led to an important transcriptional induction in the ZG in male mice, a process mirroring what is observed under HSD or LSD. This was accompanied by an activation of the Wnt/β-catenin pathway, as previously described^22^. Male mice appear to be more susceptible to DEX; possible explanations include the faster rate of adrenal cortex renewal in females and the capacity of females to replenish ZF from both subcapsular stem/precursor cells and ZG cells^18^. This would also explain why females do not increase their ZG cell pool, as seen in ST, to cope with cell loss and ZF atrophy, in contrast to what is observed in males.

Overall, our experiments also show only slight changes in cell lineage conversion following a LSD or HSD or following DEX treatment. Although this is in agreement with the lack of major structural modifications following a LSD or HSD, but not DEX treatment, one alternative explanation may be that the speed of lineage conversion in our model does not allow to well identify such changes, as transdifferentiation at the start of the treatment at 4 weeks of age has well progressed into the entire adrenal cortex. An inducible model of Cre recombination allowing to activate the recombinase at the beginning or immediately before beginning of the challenge may allow to more precisely address the effect of challenges on cell lineage conversion. Indeed, our results show that both salt diets and DEX treatment affect the regression of the ZX with time and cell lineage conversion rate from the ZF, indicating that the ZX is affected by challenges affecting the ZG and ZF, and that effects on cell lineage conversion may become evident in a slower proliferating zone.

In conclusion, our study provides new insight into adrenal cortex cell lineage conversion and its dynamic adaptation to pathophysiological challenges. We reveal a sexually dimorphic cellular and transcriptional response to salt diets and DEX treatment, involving all layers of the adrenal cortex, and the reciprocal contribution of ZG and ZF cells to this response. Future investigations may allow to further define adrenal cortex cell dynamics and cell-specific transcriptional remodelling towards adaptation to environmental cues. In addition, these data may serve as a framework to better understand how genetic susceptibility may modulate this response, eventually leading to aldosterone dysregulation and PA in extreme cases.

## Materials and methods

### Handling and generation of Cyp11b2^+/Cre^-*m*T*m*G mouse model

Animal studies were conducted according to the guidelines and regulations formulated by the European Commission for experimental use (Directive 2010/63/EU). The use of the animals and the experimental protocol were approved by the local Ethics committee of Paris Cité University (N° 17-020) and by the French Ministère de l′Enseignement Supérieur, de la Recherche et de l′Innovation (APAFIS authorization number #11509-201703281427107 v4) and were conducted in accordance with the relevant codes of practice for the care and use of animals for scientific purposes. All animals were group-housed and bred in a dedicated husbandry specific pathogen-free (SPF) facility with 12 h/12 h light-dark cycles, had ad libitum access to food and water, and were checked daily. Their health status was routinely checked to maintain SPF grade. The *Cyp11b2^Cre^* mouse model was generated and validated on a C57BL/6J congenic background by GenOway (France). To generate the tissue specific reporter mouse model, the *Cyp11b2^+/Cre^*mice were crossed with theGt(ROSA)^26Sortm4(ACTB-tdTomato-EGFP)Luo/J^ reporter mouse model (Jackson:007676), bearing a two-color fluorescent Cre-reporter allele, to generate the *Cyp11b2^Cre^*-*m*T*m*G line. In the absence of Cre recombination, tdTomato (*m*T) fluorescence is widespread in all cells/tissues. Cells expressing Cre recombinase and future cell lines derived from these cells exhibit eGFP (*m*G) expression located at the plasma membrane instead of *m*T fluorescence. All experiments were conducted on male and female heterozygous Cre mice. To respond to ethical requirement and reduce the number of animals used, the same control mice were used for different experiments: 4-weeks old mice for age studies and Ctrl T_1_ LSD, HSD and DEX, 6-weeks old mice for age studies and Ctrl T_2_ LSD and HSD, 7-weeks old mice for age studies and Ctrl T_2_ DEX. For genotyping the following primers were used: Cre Forward: GCTCCTGCTTCACCATGTGAGCAG; Cre Reverse: GG ACCAGCTATAGAGACCTCACCAACAC.

### Blood and plasma collection

Blood was collected from the retro-orbital venous plexus with heparin-coated capillaries under isoflurane (3%) anaesthesia. Blood was centrifugated at 2000 rcf for 15 minutes at 4°C to obtain plasma. Plasma was stored at -20°C for further analysis.

### Diets

Male and female mice were fed different diets for a period of two weeks, from the age of 4 weeks until reaching 6 weeks of age (Figure 2a, and 4a, time point 1, T1): Standard (0.28% Na^+^; 0.86% K^+^, A03SP-10, SAFE, France), Low salt diet, LSD (<0.03% Na^+^; 0.8% K^+^, E15430-247, Ssniff, Soest, Germany), and High salt diet, HSD (3.2% Na^+^; 0.97% K^+^) (E15433-347, Ssniff, Soest, Germany). Half of the mice were sacrificed at the end of the period, while the others returned to a standard diet and were housed under normal conditions for 1 week and sacrificed at the age of 7 weeks (time point 2, T2). For all timepoints, the corresponding control animals were investigated (Ctrl T1, Ctrl T2).

### Dexamethasone treatment

Male and female mice were subjected to dexamethasone (DEX) treatment from the age of 4 weeks for a period of two weeks until reaching the age of 6 weeks. Half of the mice were sacrificed at the end of the treatment (time point 1, T1) while the others were removed from the treatment and housed under normal conditions for 3 weeks of recovery and sacrificed at the age of 9 weeks (time point 2, T2). For both timepoints, the corresponding control animals were investigated (Ctrl T1, Ctrl T2). DEX (DEXFREE eye drops, 1mg/ml) was administered via drinking water, protected from the light, at a concentration of 0.5% (m/v) ab libitum and changed every 48 hours to avoid compound degradation. The taste of water is not altered, and the volume of water consumed by mice under treatment was comparable to that of control mice. The mice under treatment were housed in cages of four animals maximum to avoid possible difficulties of access to water.

### Western blot

6 weeks old WT, *Cyp11b2^+/Cre^* and *Cyp11b2^Cre/Cre^* female and male mice were used for western blot analysis. Total proteins were extracted using RIPA buffer (RB 4475, BioBasic Canada) with EDTA-free protease and phosphatase inhibitor (Bimake), Tissue lysates were then spinned down for 30 min at 4°C and centrifuged at 13000 rpm for 15 min at 4°C. Protein concentration was determined using Bradford protein essay (Biorad). 20 µg of total proteins were loaded on 10% Acrylamide SDS-PAGE gel, transferred onto nitrocellulose membrane and tagged with the following antibodies: mCyp11b2 (1:500 in TBS-T 5% milk, generous gift from CE Gomez-Sanchez), goat anti-rabbit IgG (H+L)-HRP conjugate antibody (1/5000, #1706515, Bio Rad), Vinculin (1/5000, V9131, Sigma-Aldrich) and anti-mouse IgG, HRP-linked antibody (1/2500, #1706516, Bio Rad). The signal was developed by Clarity Max™ Western ECL substrate (Biorad, Hercules, CA), detected by Fujifilm Las-4000 mini Luminescent image analyser (Fujifilm, Tokyo-Japan) and quantified by Multi gauge software (Fujifilm, Tokyo-Japan). The expression of total proteins was normalized to the expression of the housekeeping protein Vinculin.

### Histological examination and immunostaining

Mice were anesthetized with isoflurane (3%) and after confirming the lack of paw-pinch reflex, sacrificed by cervical dislocation after blood sampling. Right adrenal was snap frozen in liquid nitrogen and left adrenal was processed for histology as following. The tissue was fixed in 4% PFA O/N and partially dehydrated through successive immersions into 10%, 20% and 30% sucrose solutions for 24 hours each. Adrenals were subsequently placed in embedding moulds and embedded with pure optimal cutting temperature (OCT) compound and isopentane and stored at −80 °C. Adrenals were sectioned in 10 μm cuts using an Epredia™ Cryostat CryoStar™ NX70, at a temperature of -30°C, transferred to a Superfrost Plus^TM^ Microscope Slides and then stored at -80°C. For histological analysis H&S staining was performed using the Rapid Frozen Sections H&S staining kit (Bio-Optica, 04-061010) following the manufacturer’s instructions and dehydrated through 10 immersions in 95% ethanol, 10 immersions in 100% ethanol and 20 immersions in Xylene. The sections were then mounted using Pertex (Histolab, 00840). For immunofluorescence studies tissues were permeabilized for 15 min either with TBS 0.1% Triton X100 (mCyp11b2 and Ki67) or PBS 0.1% Triton X100 (20αHSD and Dab2). The sections were then blocked with 5% BSA and 10% normal donkey serum (NDS) diluted in TBS 0.05% Triton X100 (mCyp11b2, Ki67) or 10% NDS in PBS 0.1% Triton X100 (20αHSD and Dab2) for 2 hours at room temperature. For mCyp11b2 and Ki67 detection primary antibodies were diluted in 3% BSA and 3% NDS diluted in TBS 0.05% Triton X100 and incubated overnight at 4°C whereas for 20αHSD and Dab2 detection, the primary antibodies were diluted in 1% NDS in PBS 0.1% Triton X100. The following primary antibodies were used: mouse Cyp11b2 (mCyp11b2, 1/100, generous gift from CE Gomez-Sanchez), Dab2 (1/1600, Cell Signalling, D709T), Ki67 (1/1000, Abcam, ab15580), 20αHSD (1/500, generous gift from Yacob Weinstein). Donkey anti-rabbit Alexa 647, diluted 1/1000 in 3% BSA or 3% NDS diluted in PBS/TBS 0.05% Triton X100 was used as secondary antibody for 2 hours at room temperature. Nuclei were counterstained using 4′,6-diamidino-2-phenylindole (DAPI) (1/5000, Roche Diagnostics GmbH). Whole slide imaging (WSI) was performed for all the sections using the Olympus Slideview VS200 digital slide scanner (Olympus Corporation, Tokyo, Japan). The region of interests (ROIs) and focus points in the ROI were automatically recognized by the scanner and, in order to reach a high image quality, focus points were manually updated by the user. The slides were scanned using the UPlanXApo 20x/0.80 objective lens, which provides a high numerical aperture for superior resolution and image clarity.

### Image analysis

The quantitative analysis of the immunofluorescence images was performed using HALO software (version 3.6.4134, Indica Labs) with the HighPlex Fl v4.2.14 module. The fluorescence TIFF images were imported into HALO. Background correction was applied to reduce non-specific fluorescence and autofluorescence in the tissue. All three channels (DAPI, Alexa Fluor 488, Alexa Fluor 594 and Alexa Fluor647) were aligned to ensure accurate signal quantification. ROIs corresponding to the adrenal tissue were manually annotated. Artefacts, left adipose tissue around the adrenal and the medulla were manually removed and excluded from the analysis. DAPI was used for traditional nuclear segmentation within the ROIs. Alexa Fluor 488 was used to detect the GFP^+^ membrane and Alexa Fluor 594 to detect the mTomato in the Cyp11b2^+/Cre^-*m*T*m*G mice. Cyp11b2-, Dab2-, Ki67- and 20αHSD-positive cells were identified using Alexa Fluor 647. Fluorescence intensity thresholds were set for all the channels based on the intensity of the signal.

### Bi-photon imaging

The images were acquired with the Leica SP8-Dive biphoton. The tile scans were lunched with Z-stack 125um, Z step size 2um. and a 3D visualization and 3D reconstruction followed by the mosaic merge with LSA X software.

### In situ mRNA analysis

In situ mRNA analysis was performed using the RNAscope assay (ACD, Biotechne). For RNAscope experiments, frozen slides were used of *Cyp11b2*^+/Cre^-*m*T*m*G mice. RNAscope for mouse *Cyp11b2* and *Cyp11b1* was performed according to the manufacturers’ instructions using the RNAscope 2.5 HD Duplex Detection kit (ACD, Biotechne). In brief, slides were heated for 30 min at 60°C and post-fixed with 4% PFA in 1x PBS for 15 min at 4°C. Tissues were dehydrated by consecutive baths in 50%, 70% and 100% ethanol for 5 min each and then air-dried for 5 min at room temperature. Slides were then immersed in the pre-heated 1x Target Retrieval solution for 5 min and then wash with MilliQ water. After rinsing in fresh 100% ethanol the slides were air-dried. RNAscope® Protease Plus solution was applied to the slides which were then incubated for 30 mins at 40°C. The hybridisation probe mixture (*Cyp11b1* and *Cyp11b2*) was added to the sections for 2hrs at 40°C. For amplification, AMP 1 solution was added to the slide and incubated for 30min at 40°C. The same steps were repeated for 6 cycles of amplification. A signal detection step is performed by adding Fast-RED substrate/slide for 10 min at room temperature. Four additional AMP hybridisation steps were performed for the second probe followed by a green signal detection step by adding GREEN substrate for 10min at RT. The slides were then counterstained with 50% haematoxylin for 2 min at room temperature and mounted.

### RNA extraction and RT-qPCR analysis

Total RNA was isolated from adrenal using Janke and Kunkel’s Ultra-Turrax T25 (IKA technologies, Staufen DE) and TRIzol (Invitrogen, 15596018). After Deoxyribonuclease I treatment (Invitrogen, 18068-015), cDNA was generated from 0.5 µg of RNA with with iScript™ cDNA Synthesis Kit (Bio-Rad, 1708891). Duplicates of cDNA samples were then amplified on the CFX Opus 96 Real-Time PCR System using SYBR Green Supermix (Bio-Rad, 1725274). Normalization for RNA quantity and reverse transcriptase efficiency was performed against three reference genes (geometric mean of the expression of Ribosomal 18S RNA, β2-microglobulin and Ubiquitin C). Quantification was performed using the 2^^-ΔΔCt^ analysis. Sequences of primers are listed in following table.

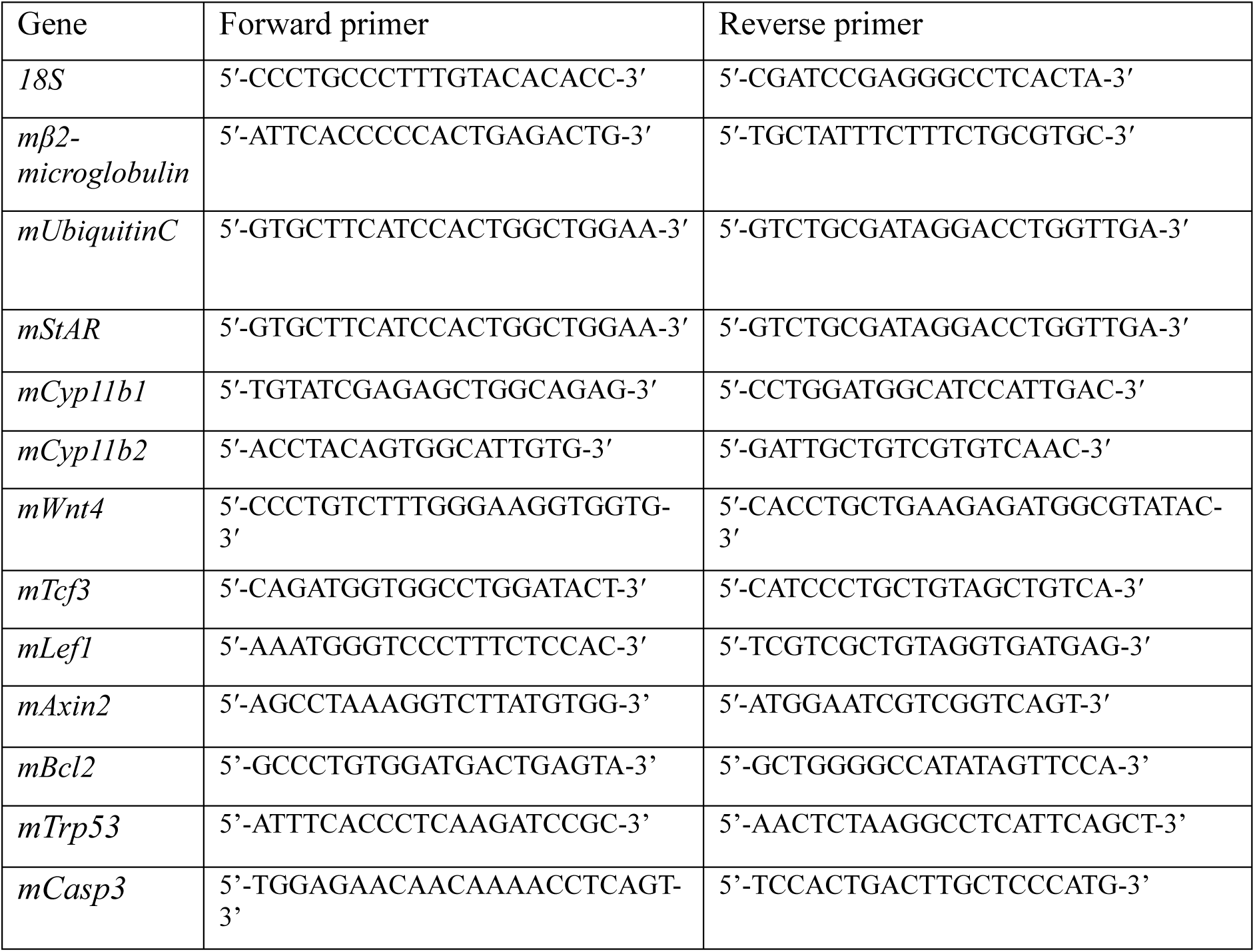

### Steroid Profiling and analysis

Plasma steroids (pregnenolone, progesterone, 11-deoxycorticosterone, corticosterone, 18-hydroxy-11-deoxycorticosterone, 18-hydroxycorticosterone, aldosterone, 18-hydroxycortisol) were simultaneously measured in a 13-minute run by Liquid Chromatography coupled to tandem Mass Spectrometry as previously described^33^. Heatmap analysis of steroid hormone concentrations was based on a Z-score normalized matrix of hormone variables, using the *pheatmap* package (v1.0.12) in R, with hierarchical clustering applied to the variables. Z-scores were calculated by subtracting the mean of each variable from individual data points and dividing by the corresponding standard deviation. A diverging color scale was used to represent the Z-score normalized hormone concentrations, ranging from deep blue (Z ≤ –1), through white (Z = 0), to deep red (Z ≥ +1). Values outside this range were capped and mapped to the deepest blue or red, to minimize the influence of extreme values on the color gradient. Principal component analysis (PCA) was performed on the original data using the *FactoMineR* package (v2.11) with default standardization, and visualized with the *factoextra* package (v1.0.7). All analyses were performed in R (v4.4.2).

### Spatial transcriptomics

#### Sample Preparation

34 tissue sections (17 males, 17 females) were used for ST. Fluorescent images were acquired on a Vectra 3 Automated Quantitative Pathology Imaging System (Akoya Bioscience) and stitched using Halo software (Excilone). The spatial gene expression process including probe hybridization, probe ligation, enabled probe release with CytAssist and extension was performed using Visium CytAssist Spatial Gene Expression for FFPE, Mouse Transcriptome, 6.5mm, 4 rxns (1000521, 10X Genomics) according to the manufacturer’s instructions. Briefly, sections were decrosslinked at 95°C for 60min, then Mouse WT pairs of probes, targeting 21000 genes, were incubated overnight. Paired probes were linked then transferred on the Visium slide using the Cytassist system. The spatial gene libraries were constructed using Probe-based Library Construction (10X Genomics). Library quantification and quality assessment was performed using Qubit fluorometric assay (Invitrogen) with dsDNA HS (High Sensitivity) Assay Kit, Bioanalyzer Agilent 2100 using a High Sensitivity DNA chip and KAPA Library Quantification Kit Illumina Platforms (Roche). Libraries were then sequenced on a NovaSeq 6000 (Illumina) using paired-end 28 x 50 bp with a depth of sequencing of 25,000 reads per spot.

#### Data analysis

Sequencing reads were demultiplexed and aligned to the mouse reference transcriptome (mm10-2020-A), using the spaceranger Pipeline (v2.0.0) in default mode. Loupe browser from 10X Genomics was used to align the barcoded spot patterns, perform spots selection on the tissue and do the pathological annotation. SpotClean (v1.4.1) was used to calculate the peer-spot contamination rates and to decontaminate the gene expression profiles from the spot swapping effect. The corrected UMI counts were loaded into Seurat (v5.10) for quality control, data integration and downstream analyses. Empty spots (nCount < 1) were removed and the data from each sample were log normalised and the effect of different number of counts (nCount) among samples was removed during scaling. For each sample, the top 3000 highly variable genes were selected using the FindVariableFeatures function. Batch effect between samples was corrected using Seurat’s FindIntegratedAnchors, and the samples were integrated using cca reduction. On the integrated dataset, dimensionality reduction was performed using principal component (PC) analysis computed on the first 30 PC. FindNeighbors was used to compute the k-nearest neighbor graph on the low-dimensional embedding, and FindClusters was used to cluster the spots with multiple resolutions (from 0.8 to 2). Finally, the maximum resolution (r=2) was considered to better align with the expected cell populations, and it was used for visualisation and all downstream analyses. Cell type labels were assigned to resulting clusters based on a manually curated list of marker genes as well as previously defined signatures of the well-known adrenal gland subtypes (Table S1). All clusters were annotated, and 6589 spots were kept for further analysis.

Differential expression analysis was performed on log-normalized non-batch-corrected data using Wilcoxon testing with Bonferroni correction as implemented in the FindMarkers function. Only genes with adjusted p-values < 0.05 were selected as significant. The lists of differentially expressed genes were further divided into UP and DOWN regulated genes based on the avg_log2FC; avg_log2FC >0 for the UP regulated genes and 4avg_log2FC <0 for the DOWN regulated ones. The signature scores were calculated using the function AddModuleScore_UCell from UCell, and a combination of Scpubr (v2.0.2) and ggplot 2 was used in all the figures. GSEA analysis was performed using the fgsea package (v1.28.0) and the Kegg gene sets belonging to the Mouse MSigDB Collections (v2024.1). As each spot might contain a mixture of cell populations, deconvolution analysis using a reference-free approach was required to recover the correct organisation of the tissues. The deconvolution of the spatial spots was performed using the Stdeconvolve [] package. The spot coordinates, the raw counts and the cell annotation were extracted directly from the Seurat object and the pixels with less than 100 genes and 10 counts were removed. The deconvolution was performed assuming a maximum of 9 cell populations and the parameters that led to the minimum perplexity of the LDA model were selected for downstream analysis. The pie charts and all the figures were generated using a combination of ggplot and the internal function vizAllTopics^34^. In total, we deconvolved 4 main populations, ZG, ZF, ZX and Medulla, with additional components associated with a mixture of the main populations and immune cells (Figure S17). We then compared the result of the deconvolution and the spatial expression patterns of the markers used for the annotation. In both cases, there was a good correlation with the spatial organisation observed in the histological images. In particular, the regions with a high degree of mixture were found either in the peripheral zones between distinct layers of the tissue or in sections where the tissues were folded and overlapped (Figure S17).

### Statistical analyses

Each symbol in graphical representations show data Plot show mean ± SEM Quantitative variables are reported as means ± SEM when Gaussian distribution or medians and interquartile range when no Gaussian distribution. Parametric tests were applied only if normal distribution could be confirmed. Pairwise comparisons were done with unpaired t-test or Mann-Whitney test respectively; global comparison was evaluated using ANOVA or Kruskall Wallis test, followed by Sidak’s or Dunn’s multiple comparison test. A p value < 0.05 was considered significant for comparisons between 2 groups. Analyses were performed using Graphpad Prism 10 (GraphPad software Inc, San Diego, CA).

## Supporting information

Faedda et al_Supplemenantary material

## Acknowledgments

This work was funded through the European Union’s Horizon 2020 research and innovation programme under the Marie Sklodowska-Curie grant agreement No. 954798 (MINDSHIFT804 ITN) institutional support from INSERM, by the Agence Nationale pour la Recherche (ANR-18-CE93-0003-01), the Fondation pour la Recherche Médicale (EQU201903007864).

## Notes

### Competing Interest Statement

The authors have declared no competing interest.

