## Supplementary material for "Sexually dimorphic interzonal crosstalk reshapes the adrenal cortex in response to pathophysiological challenges": Faedda et al_Supplemenantary material

### Supplementary Figures and Tables

#### Figure S1

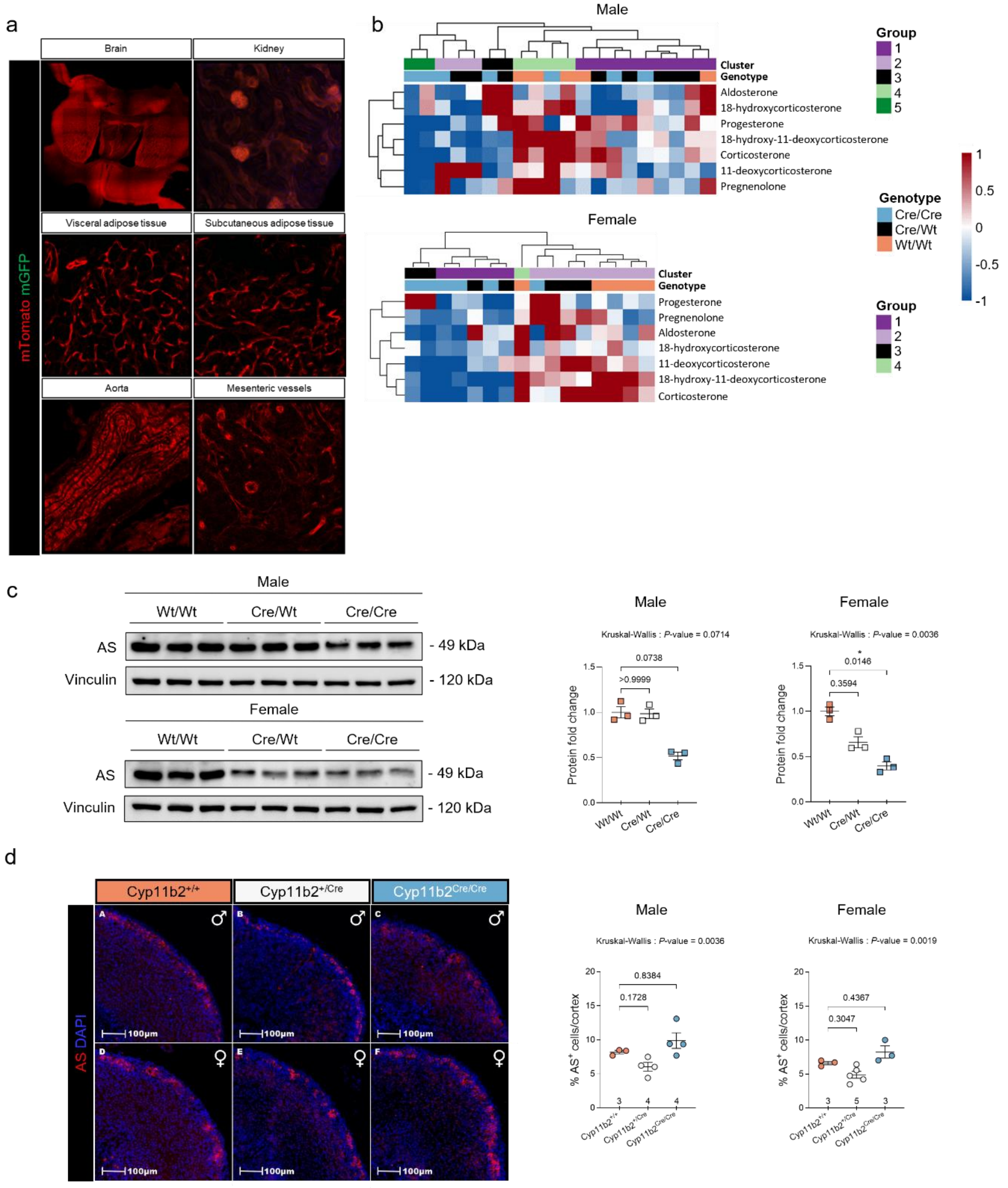


**Figure S1. Characterization of the adrenal phenotype of the *Cyp11b2*$^{Cre}$ mouse model. (a).** Analysis of mGFP and mTomato expression in the brain, kidney, visceral and subcutaneous adipose tissue, aorta and mesenteric vessels of a 5-weeks old *Cyp11b2*$^{+/Cre}$-*m*T*m*G male mouse **(b).** Hierarchical clustering heat map visualization of plasma steroid profiles of *Cyp11b2*$^{+/+}$ (n=5), *Cyp11b2*$^{+/Cre}$ (n=8) and *Cyp11b2*$^{Cre/Cre}$ (n=7) male (upper pannel) and of *Cyp11b2*$^{+/+}$

(n=5), *Cyp11b2*$^{+/Cre}$ (n=5) and *Cyp11b2*$^{Cre/Cre}$ (n=6) female (lower pannel) mice of 6 weeks of age measured by LC-MS/MS. **(c).** Representative immunoblots and histograms showing Adosterone Synthase (AS) expression in wild type *Cyp11b2*$^{+/+}$, heterozygous *Cyp11b2*$^{+/Cre}$ and homozygous *Cyp11b2*$^{Cre/Cre}$ 6 weeks old mice of both sexes, relative to Vinculin. P values were obtained with Kruskal-Wallis test and adjusted with Dunn's multiple comparisons test. **(d).** Immunofluorescent labelling of AS (red) on adrenals from wild type *Cyp11b2*$^{+/+}$, heterozygous *Cyp11b2*$^{+/Cre}$ and homozygous *Cyp11b2*$^{Cre/Cre}$ 6 weeks old male and female mice. P values were obtained with Kruskal-Wallis test. Data are presented as mean values ± SEM. Data are presented as mean values +/- SEM. *P value < 0.05; **P value < 0.01; ***P value < 0.001.

Figure S2

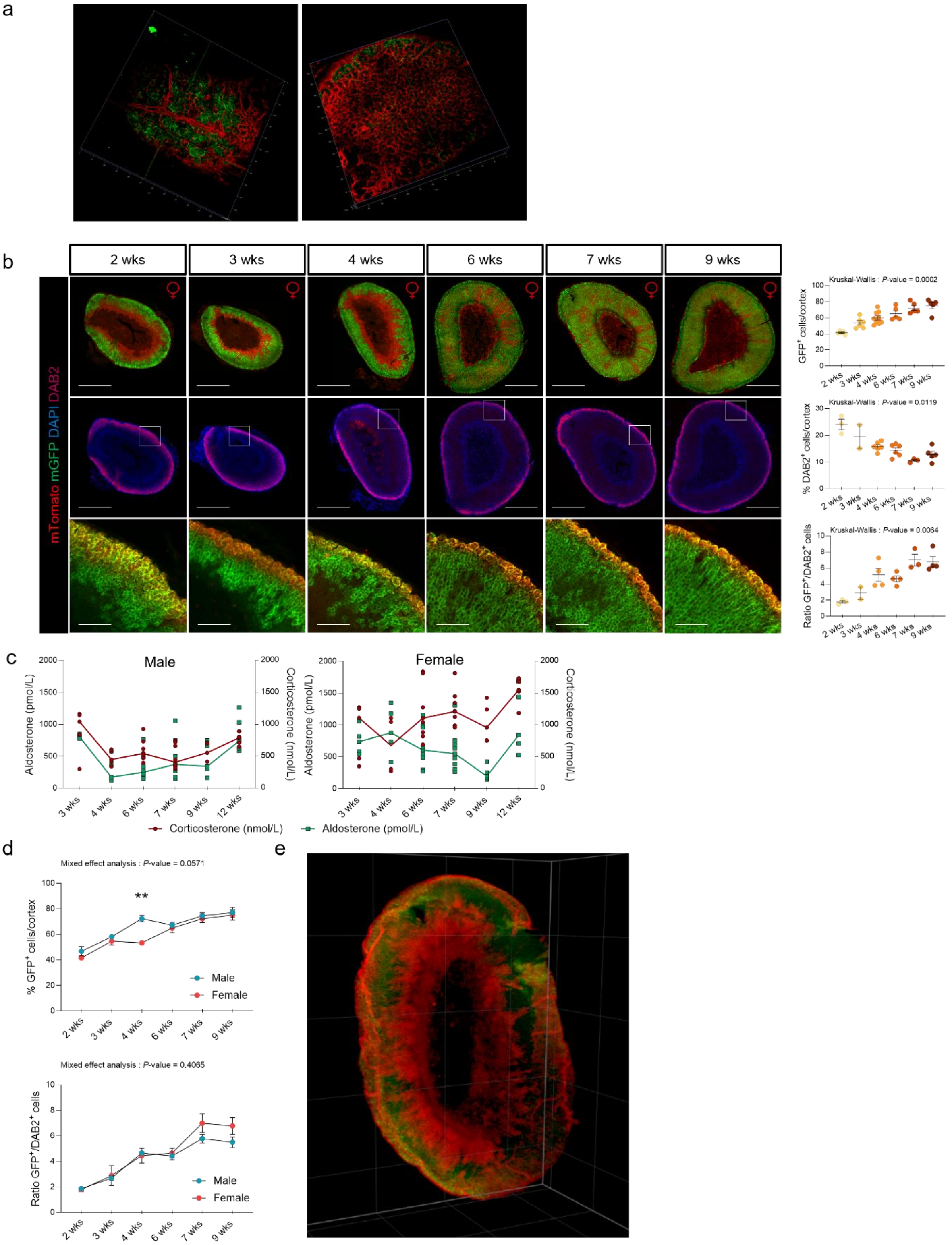


**Figure S2. Transdifferentiation kinetics in *Cyp11b2*$^{Cre}$::*m*T*m*G mice. (a).** Biphoton 3-D images of mGFP and mTomato expression of *Cyp11b2*$^{+/Cre}$-*m*T*m*G adrenals from a male mouse at postnatal day 1 (P1) after clarification; images represent superficial (left) and median (right) z-stacks of the adrenal. Specific mGFP is localized to the membrane, while unspecific background signal is localized to the cytoplasm. **(b).** Coimmunofluorescence of mGFP (green) with Tomato (red) and DAB2 (ruby red) of *Cyp11b2*$^{+/Cre}$-*m*T*m*G female mice adrenals at 2, 3, 4, 6, 7 and 9 weeks of age and quantification of the DAB2$^{+}$ cells, mGPF$^{+}$ cells and analysis of cell lineage conversion expressed as the ratio of the number of cells expressing mGFP over the cortex area or the number of cells expressing DAB2. P values were obtained with Kruskal-Wallis test and adjusted with Dunn's multiple comparisons test. Scale Bar: 500 µm (upper panels),100 µm (lower pannels). White squares represent the magnification area in the lower panel. **(c).** Aldosterone and corticosterone plasma levels in male (left panel) and female (right panel) *Cyp11b2*$^{+/Cre}$-*m*T*m*G mice from 2 to 12 weeks of age. **(d).** Mixed effect analysis of cell lineage conversion by quantification of mGFP represented as %GFP$^{+}$ cells/cortex (upper panel) or %GFP$^{+}$ cells/%Dab2 $^{+}$ cells (lower panel) in male and female *Cyp11b2*$^{+/Cre}$ –*m*T*m*G mice at 2, 3, 4, 6, 7 and 9 weeks of age. Statistical analysis was performed using two-way analysis and adjusted with Šídák's multiple comparisons test. **(e)**. Volume rendering of a wholemount lightsheet fluorescence microscopy (LSFM) imaged, EZ Clear processed *Cyp11b2*$^{+/Cre}$-*m*T*m*G female 6 weeks old mouse adrenal tissue digitally sectioned along the longitudinal axis using Slide Book digital microscopy software. Data are shown as mean ± SEM. *P value < 0.05; **P value < 0.01; ***P value < 0.001.

**Figure S3**

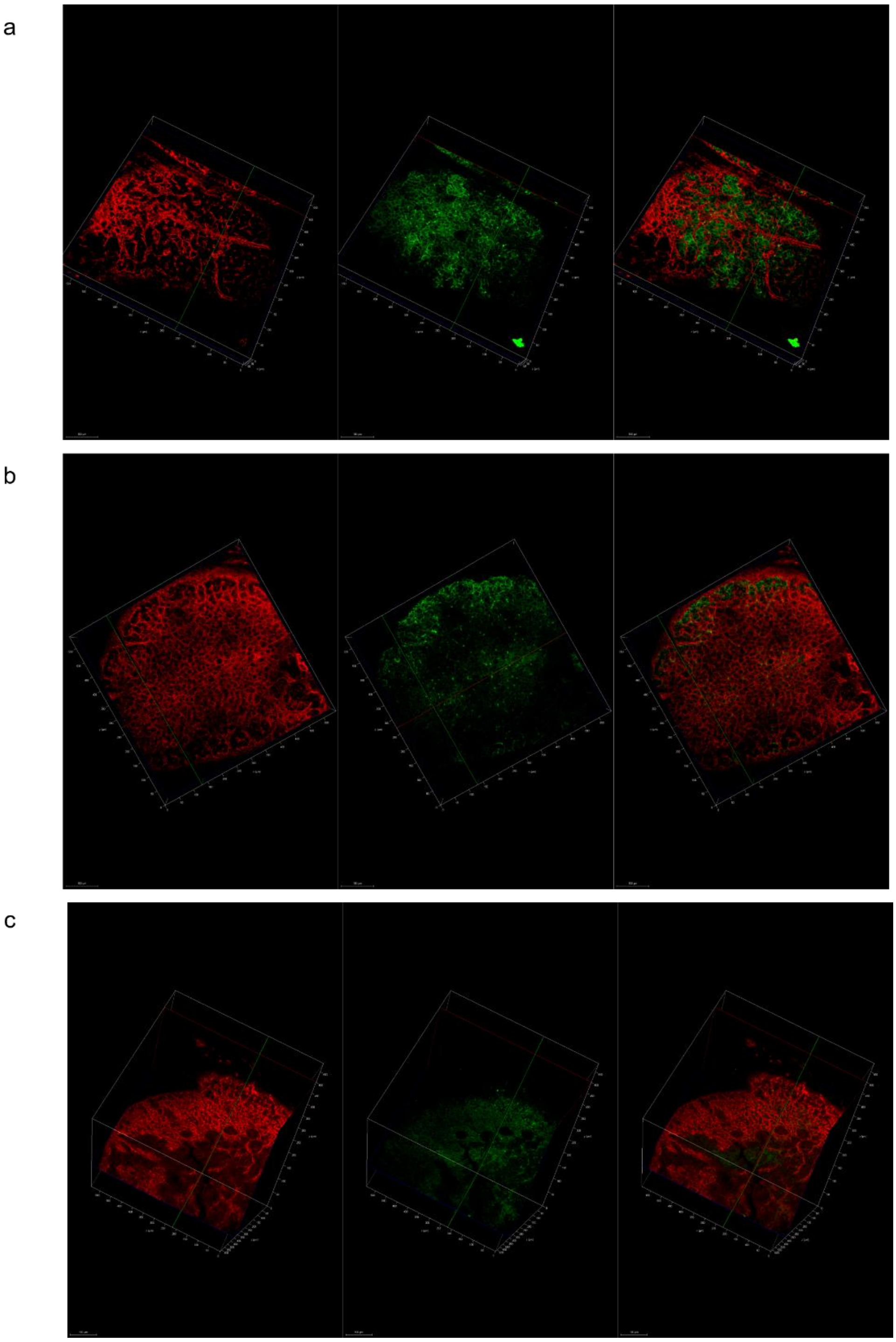

**Figure S3. Transdifferentiation in *Cyp11b2*$^{Cre}$::*m*T*m*G mice at P1**. **(a-c).** Biphoton 3D images of *Cyp11b2*$^{+/Cre}$-*m*T*m*G adrenals after clarification at postnatal day 1 (P1), with mTomato (left panel), mGFP (middle) and overlay (right panel). **(a).** Superficial z-stack of a *Cyp11b2*$^{+/Cre}$-*m*T*m*G mouse. **(b).** Median z-stack of a *Cyp11b2*$^{+/Cre}$-*m*T*m*G mouse. **(c).** Biphoton 3D images of *Cyp11b2*$^{+/+}$-*m*T*m*G control adrenal. Specific mGFP is localized to the membrane, while unspecific background signal is localized to the cytoplasm.

### Figure S4

a

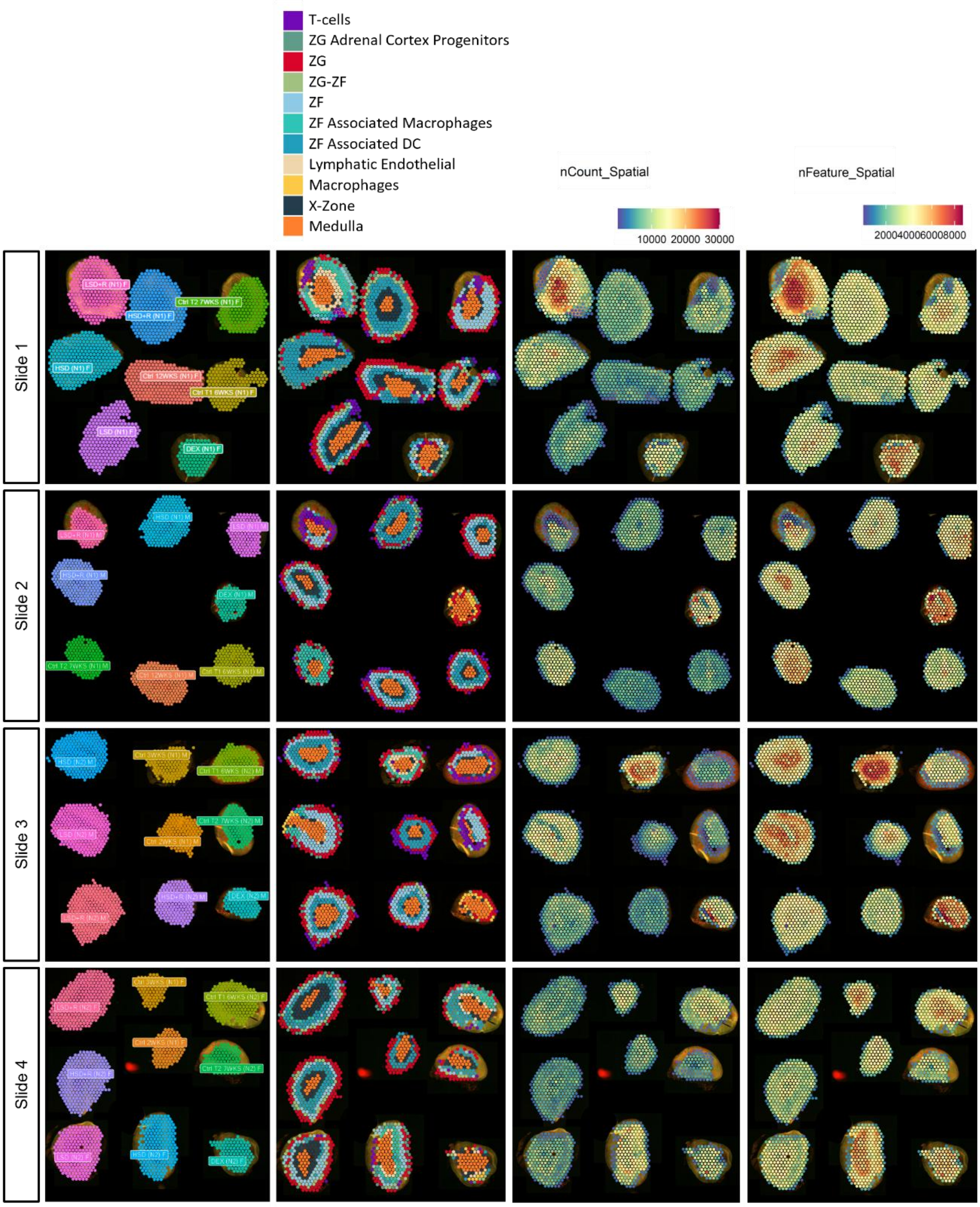


**Figure S4. Spatial transcriptomic quality control. (a)** Spatial arrangement of the different adrenal slices of control and treated mice and localization of the 11 cell clusters on the four 10x Visium slides with the number of unique transcripts (nCount) or unique genes (nFeature) per spatial spot.

**Figure S5**

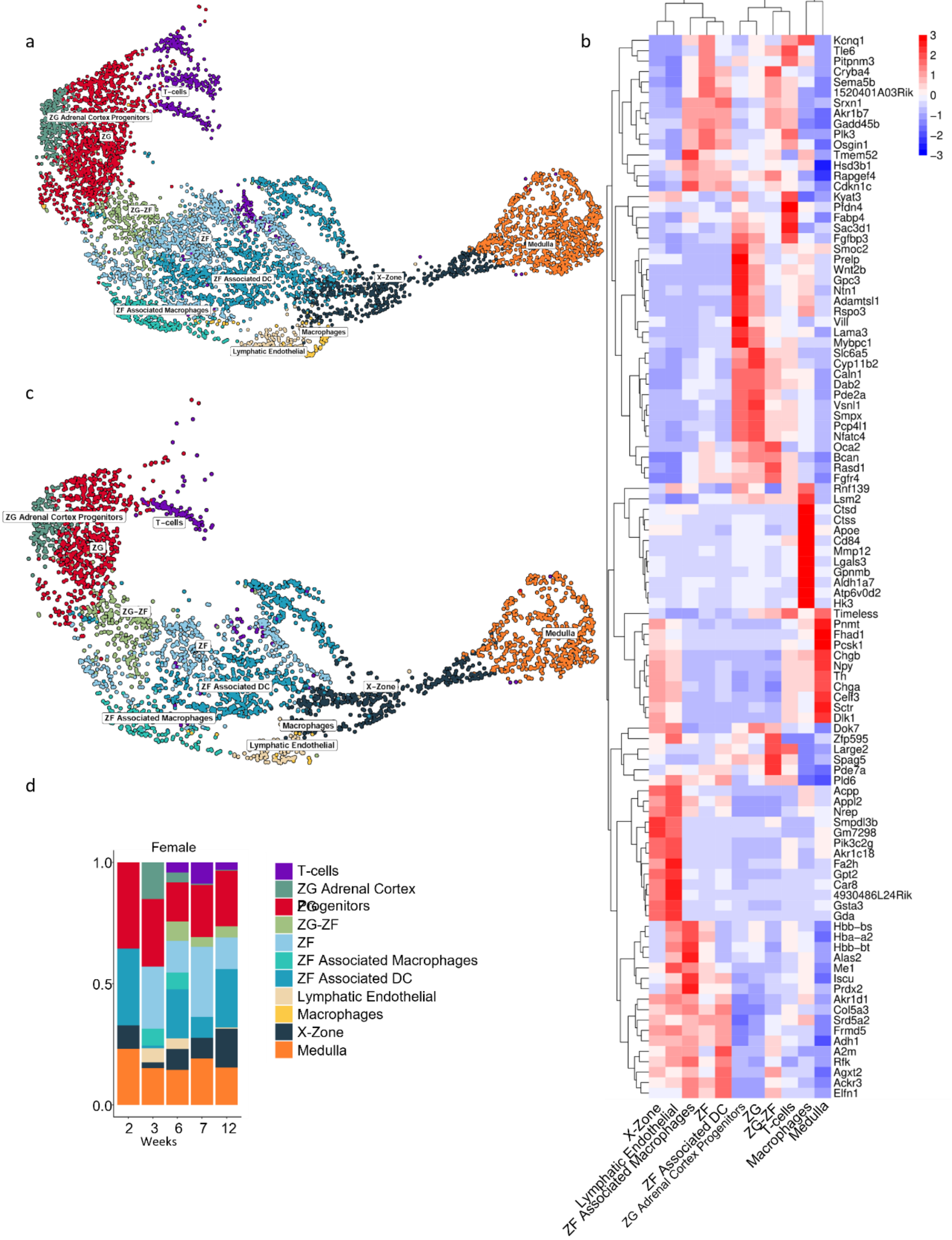


**Figure S5. Adrenal spatial transcriptomic analysis at different ages. (a)**. General uMAP of the spots detected by the 4 10x Visium slides, all conditions and **(b).** Heatmap of the top differentially expressed genes (DEGs) in each of the different identified cell clusters. **(c)**. UMAP of adrenal cell clusters in control female *Cyp11b2*$^{+/Cre}$-*m*T*m*G mice of 2, 3, 6, 7 and 12 weeks of age. **(d).** Cell cluster proportion of control female *Cyp11b2*$^{+/Cre}$-*m*T*m*G mice of 2, 3, 6, 7 and 12 weeks of age.

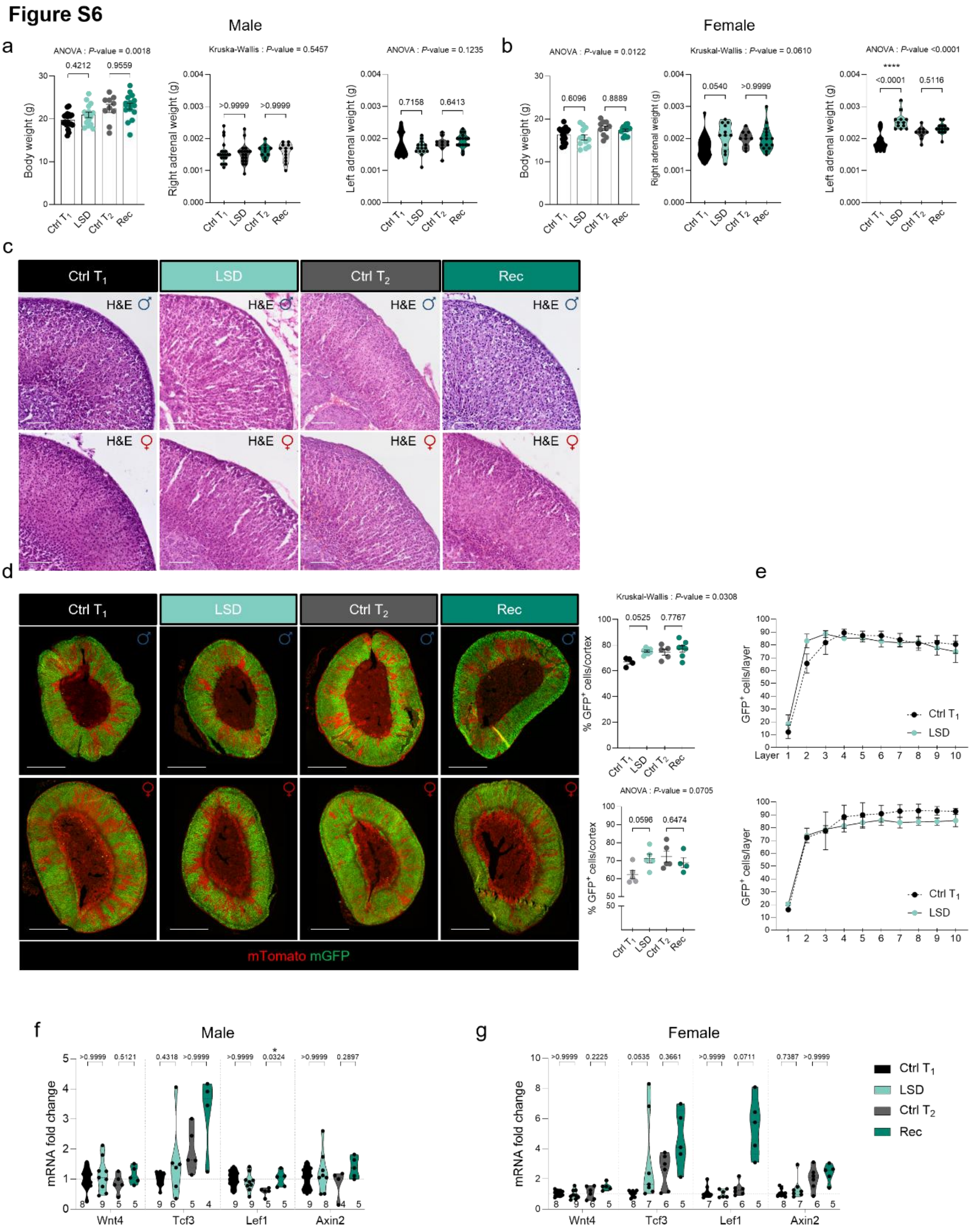


**Figure S6. Effect of LSD on the adrenal cortex. (a-b).** Body weight and absolute right and left adrenal mass of *Cyp11b2*$^{+/Cre}$-*m*T*m*G male **(a)** and female **(b)** mice fed with low salt diet from 4 to 6 weeks of age ($T_1$) and after a recovery period from 6 weeks to 7 weeks ($T_2$). **(c).** H&E staining of male and female, control and treated, *Cyp11b2*$^{+/Cre}$-*m*T*m*G mouse adrenals at $T_1$ and $T_2$. **(d).** Coimmunofluorescence of mGFP (green) with mTomato (red) and quantification of the mGPF$^+$ cells in the adrenal cortex of *Cyp11b2*$^{+/Cre}$-*m*T*m*G male (upper panel) and female (lower panel) mice at $T_1$ and $T_2$. Scale Bar: 500 µm. **(e).** Analysis of the progression of mGFP+ cells along the adrenal cortex in the first 10 layers of *Cyp11b2*$^{+/Cre}$-*m*T*m*G male (upper panel, 10 layer) and female (lower pannel, 10 layers) mice fed with low salt diet from 4 to 6 weeks of age ($T_1$) and after a recovery period from 6 weeks to 7 weeks ($T_2$). Each layer has a thickness of 25 micron. **(f-g).** Effect of LSD on the expression of Wnt/β-catenin genes in male **(f)** and female **(g)** mice. P values were obtained with Ordinary one-way ANOVA, adjusted with Šídák's multiple comparisons test or with Kruskal-Wallis test and adjusted with Dunn's multiple comparisons test, as indicated. Data are presented as mean values ± SEM. *P value < 0.05; **P value < 0.01; ***P value < 0.001.

**Figure S7**

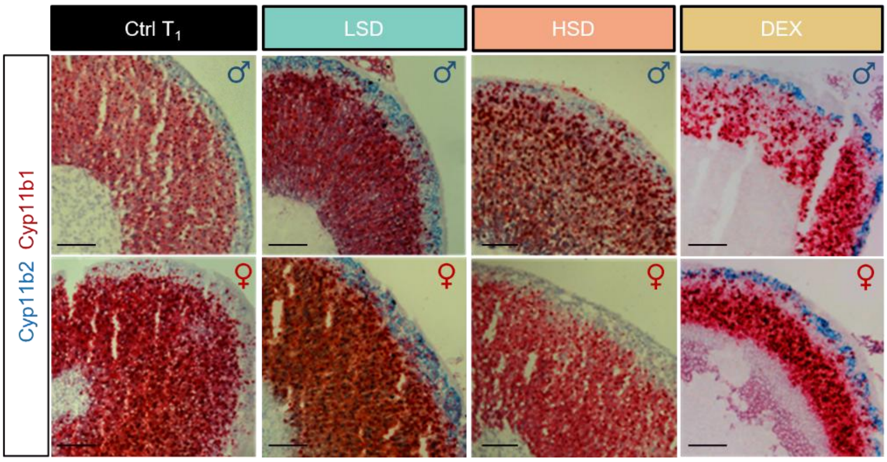


**Figure S7. Effect of LSD, HSD and DXM treatment on Cyp11b2 and Cyp11b1 mRNA expression in the adrenal cortex.** RNAscope in situ hybridization performed using one probe for *Cyp11b2* (blue) and one for *Cyp11b1* (red) in male and female mice fed a low salt (LSD) or high diet (HSD) for a two-week period, or submitted to DEX treatment. Scale Bar: 500 µm.

**Figure S8**

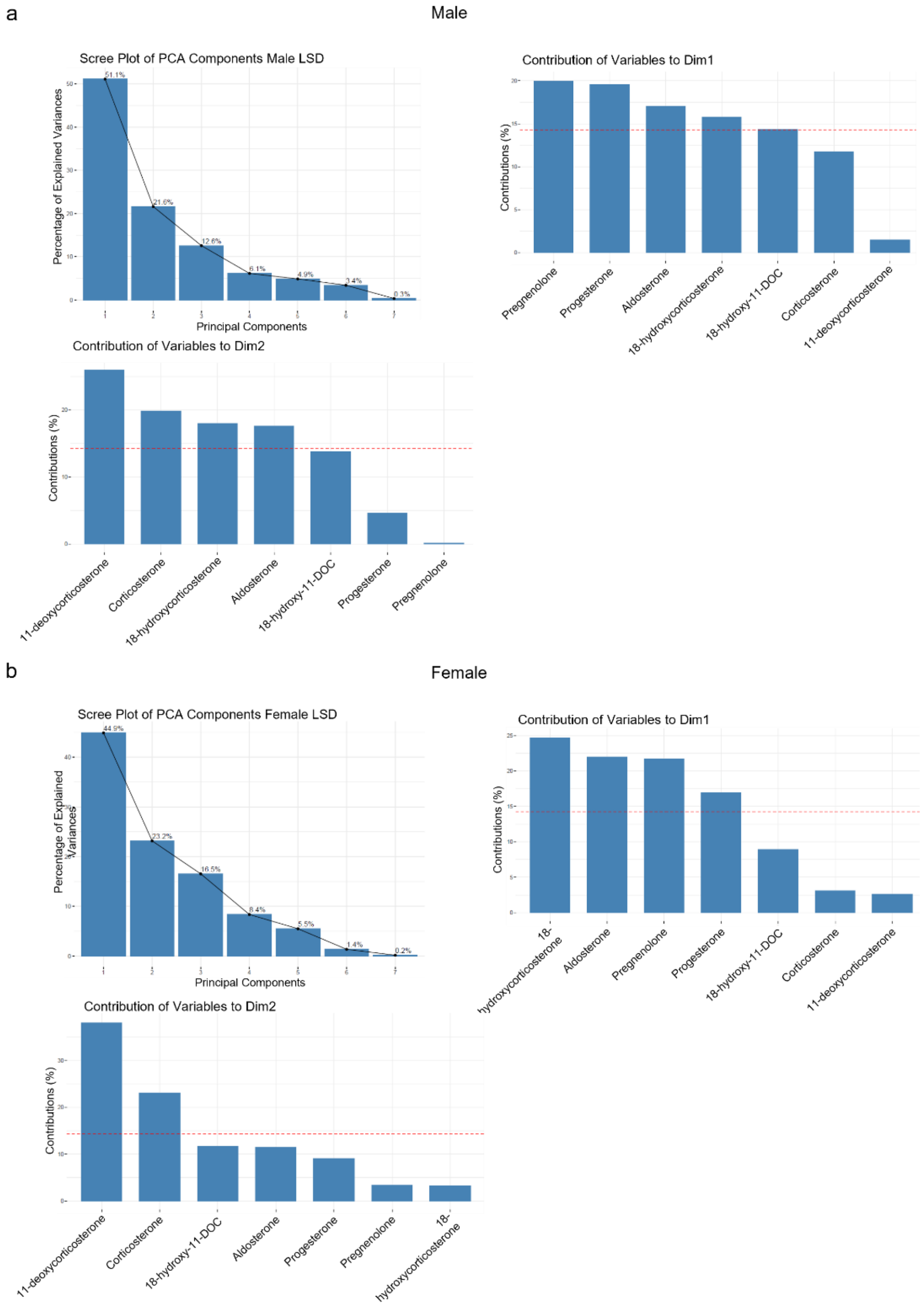


**Figure S8. Characteristics of Principal component analysis (PCA) of adrenal plasma steroid profiles following a LSD. (a).** Scree plot in which the x-axis shows the principal components (dimensions), which in this case are 7, and their contribution to variance of the Dim1 and Dim2 in male *Cyp11b2*$^{+/Cre}$-*m*T*m*G mice fed with LSD at $T_1$ and $T_2$ vs control littermates. **(b).** Scree plot in which the x-axis shows the principal components (dimensions) and their contribution to variance of Dim1 e Dim2 in female *Cyp11b2*$^{+/Cre}$-*m*T*m*G mice fed with LSD at $T_1$ and $T_2$ vs control littermates. PCA of plasma steroids revealed clear separation between control and LSD-fed mice along the first Dimension (Dim1) explaining 51.1% in males and 44.9% in females of the total variance.

**Figure S9**

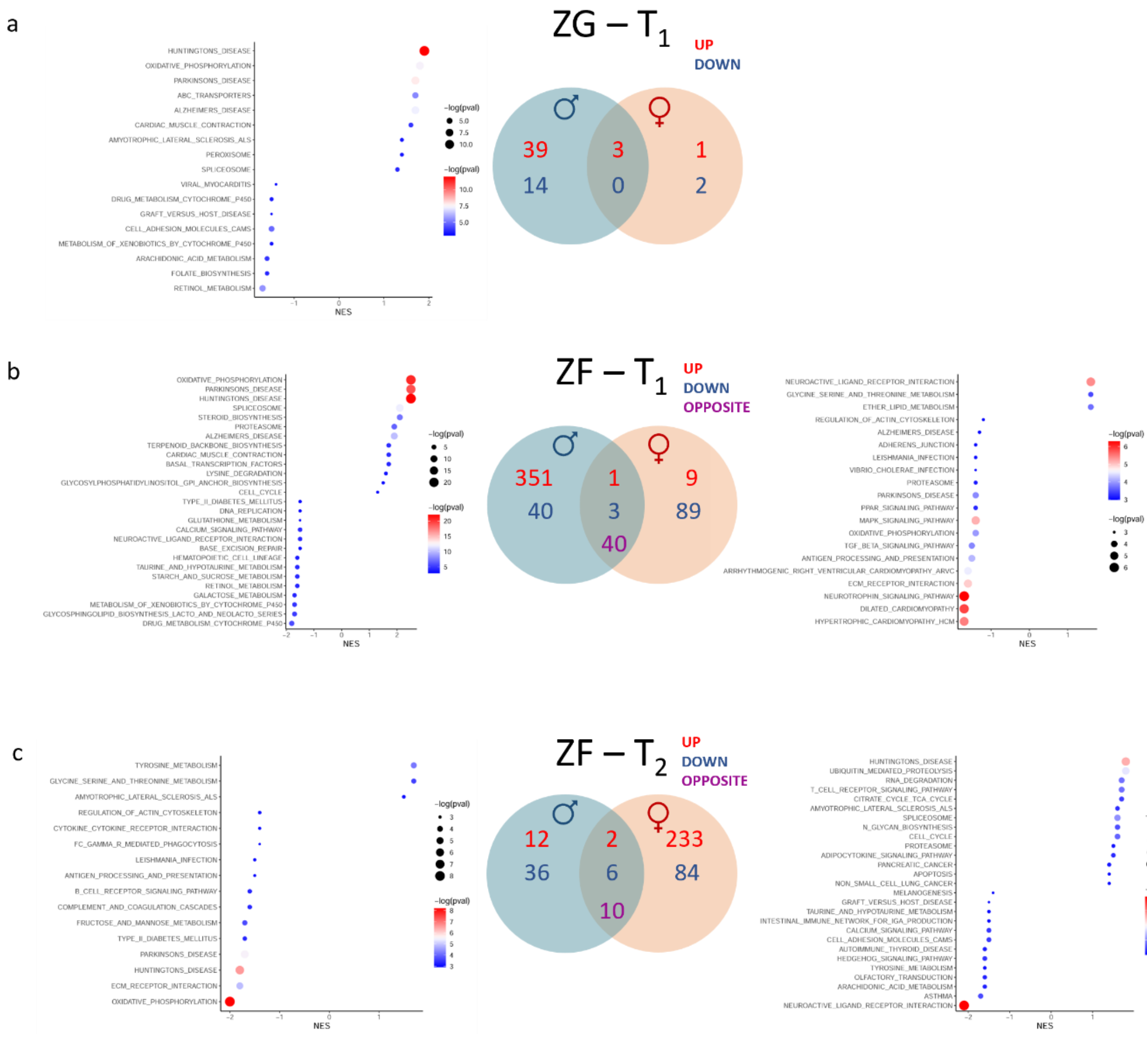


**Figure S9. Effect of LSD on zone-specific gene expression. (a).** Gene set enrichment analysis (GSEA) pathways from the KEGG database (adapted on mouse) of male *Cyp11b2*$^{+/Cre}$-*m*T*m*G LSD-treated mice vs Ctrl $T_1$ in the ZG cluster and Venn diagram illustrating the number of up and downregulated genes in male and female *Cyp11b2*$^{+/Cre}$-*m*T*m*G mice treated with LSD vs Ctrl $T_1$. **(b).** GSEA pathways from the KEGG database (adapted on mouse) of male (left panel) and female (right panel) *Cyp11b2*$^{+/Cre}$-*m*T*m*G LSD-treated mice vs Ctrl $T_1$ in the ZF cluster and Venn diagram illustrating the up and down and opposite regulated genes between males and females. **(c).** GSEA pathways from the KEGG database (adapted on mouse) of *Cyp11b2*$^{+/Cre}$-*m*T*m*G Recovery vs Ctrl $T_2$ male (left panel) and female (right panel) mice in the ZF cluster and Venn diagram illustrating the up and down and opposite regulated genes between males and females. Pathways were ordered by the NES (normalized enriched score) and only the top 20 up and down regulated pathways are visualized using dotplots. The color and size of the dots reflect the significance of the enrichment [-log(pval)].

### Figure S10

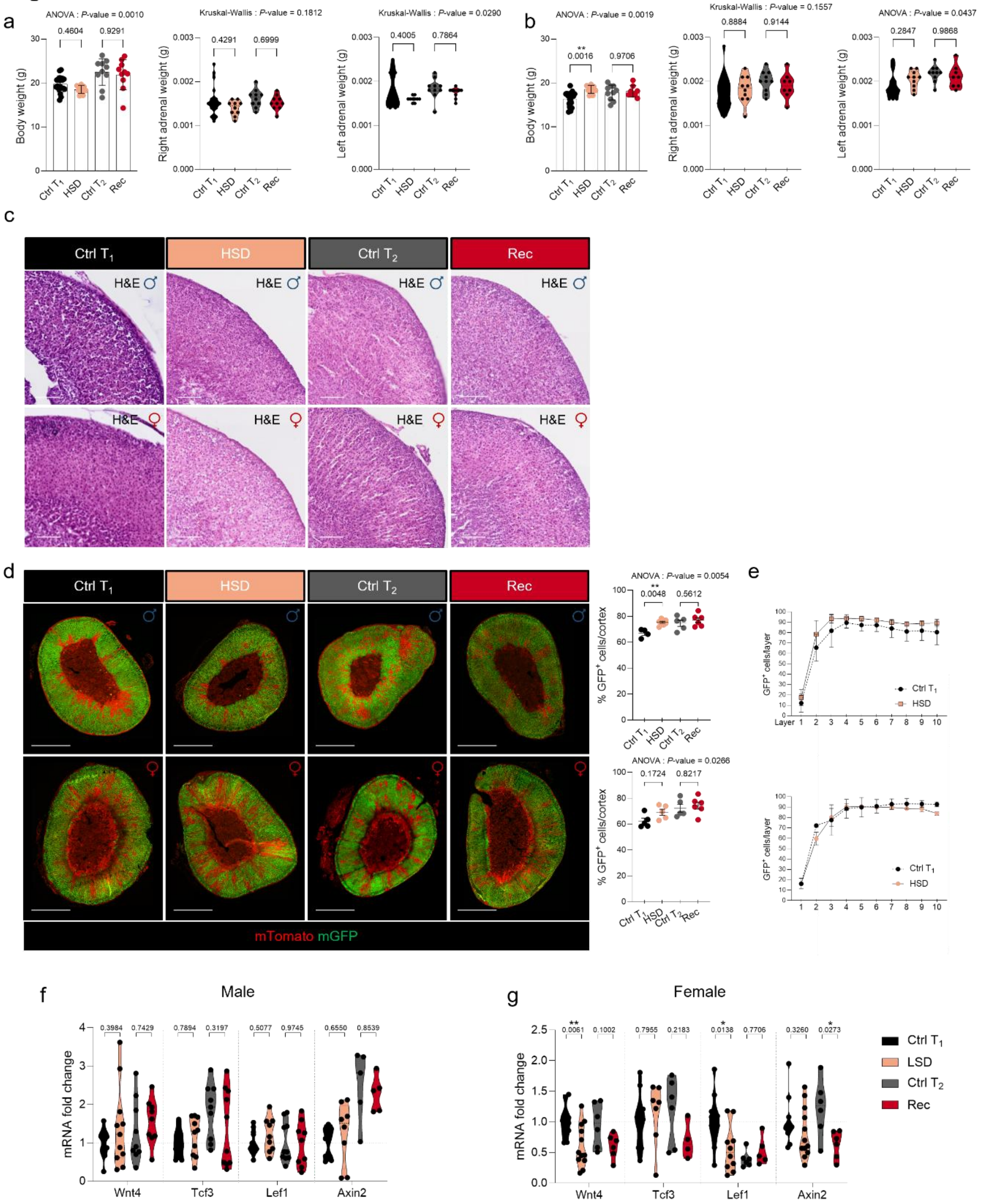

**Figure S10. Effect of HSD on the adrenal cortex. (a, b)** Body weight and absolute right and left adrenal weight of *Cyp11b2*$^{+/Cre}$-*m*T*m*G male **(a)** and female **(b)** mice fed with high salt diet from 4 to 6 weeks of age ($T_1$) and after a recovery period from 6 weeks to 7 weeks ($T_2$). **(c).** H&E staining of male (upper panel) and female (lower panel), control and treated, *Cyp11b2*$^{+/Cre}$-*m*T*m*G mouse adrenals at $T_1$ and $T_2$. **(d).** Coimmunofluorescence of mGFP (green) with Tomato (red) and quantification of the mGPF$^+$ cells in the adrenal cortex of *Cyp11b2*$^{+/Cre}$-*m*T*m*G male (upper panel) and female (lower panel) mice at $T_1$ and $T_2$. Scale Bar: 500 µm. **(e).** Analysis of the progression of mGFP+ cells along the adrenal cortex layers of *Cyp11b2*$^{+/Cre}$-*m*T*m*G male (upper panel, 10 layer) and female (lower pannel, 10 layers) mice fed with high salt diet from 4 to 6 weeks of age ($T_1$) and after a recovery period from 6 weeks to 7 weeks ($T_2$). Each layer has a thickness of 25 micron. **(f-g).** Effect of HSD on the expression of Wnt/β-catenin genes in male **(f)** and female **(g)** mice. P values were obtained with Ordinary one-way ANOVA, adjusted with Šídák's multiple comparisons test or with Kruskal-Wallis test and adjusted with Dunn's multiple comparisons test, as indicated. Data are presented as mean values ± SEM. *P value < 0.05; **P value < 0.01; ***P value < 0.001.

**Figure S11**

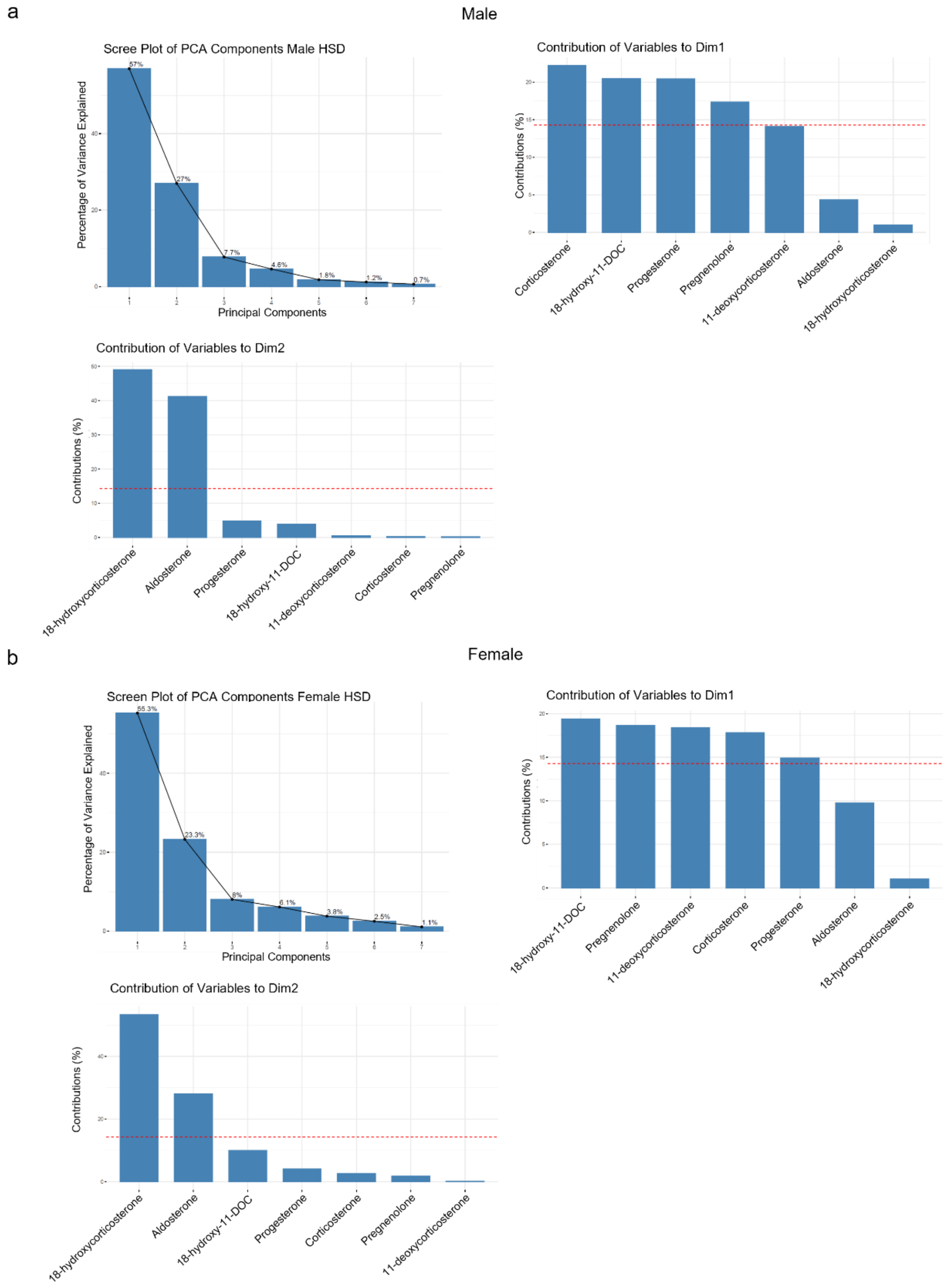


**Figure S11. Characteristics of Principal component analysis (PCA) of adrenal plasma steroid profiles following a HSD. (a-b).** Scree plot representing the principal components in relation to the percentage of explained variance and the respective contribution of single variables to dimensions 1 and 2 in male **(a)** and female **(b)** *Cyp11b2*$^{+/Cre}$-*m*T*m*G mice fed a HSD vs control littermates.

**Figure S12**

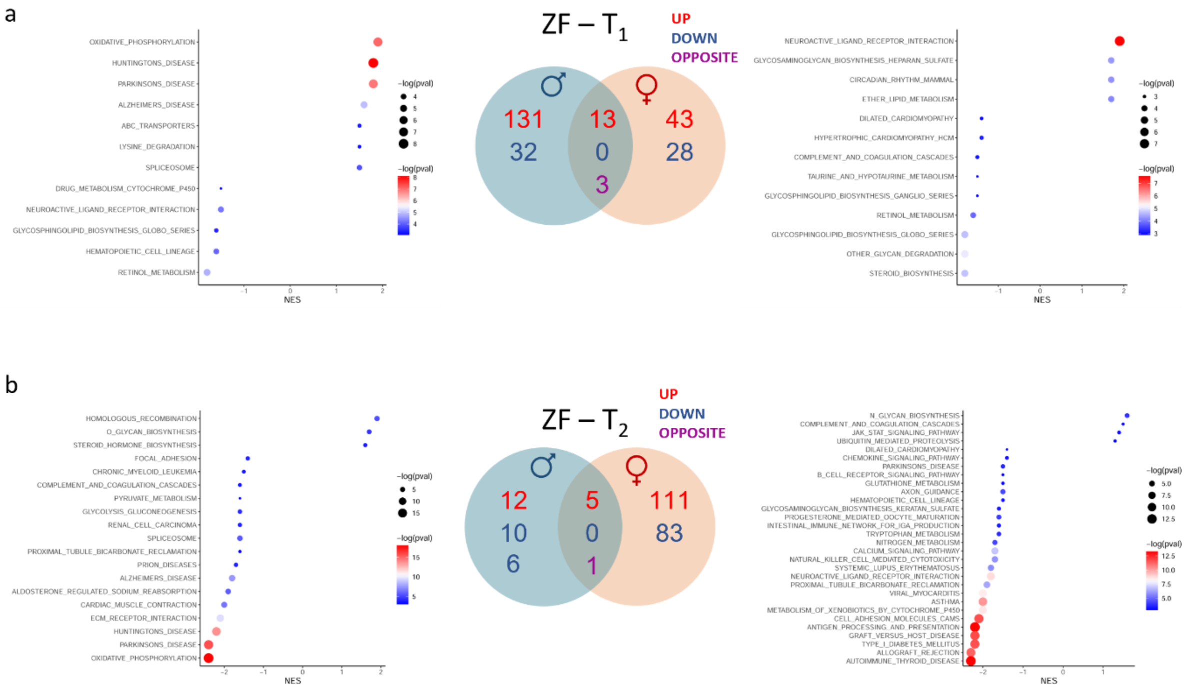


**Figure S12. Effect of HSD on zone-specific gene expression. (a).** Gene set enrichment analysis (GSEA) pathways from the KEGG database (adapted on mouse) of male (left panel) and female (right panel) *Cyp11b2*$^{+/Cre}$-*m*T*m*G HSD-treated mice vs Ctrl $T_1$ in the ZF cluster and Venn diagram illustrating the up and down and opposite regulated genes between males and females. **(b).** GSEA pathways from the KEGG database (adapted on mouse) of *Cyp11b2*$^{+/Cre}$-*m*T*m*G Rec vs Ctrl $T_2$ male (left panel) and female (right panel) mice in the ZF cluster and Venn diagram illustrating the up and down and opposite regulated genes between males and females. Pathways were ordered by the NES (normalized enriched score) and only the top 20 up and down regulated pathways are visualized using dotplots. The colour and size of the dots reflect the significance of the enrichment [-log(pval)].

### Figure S13

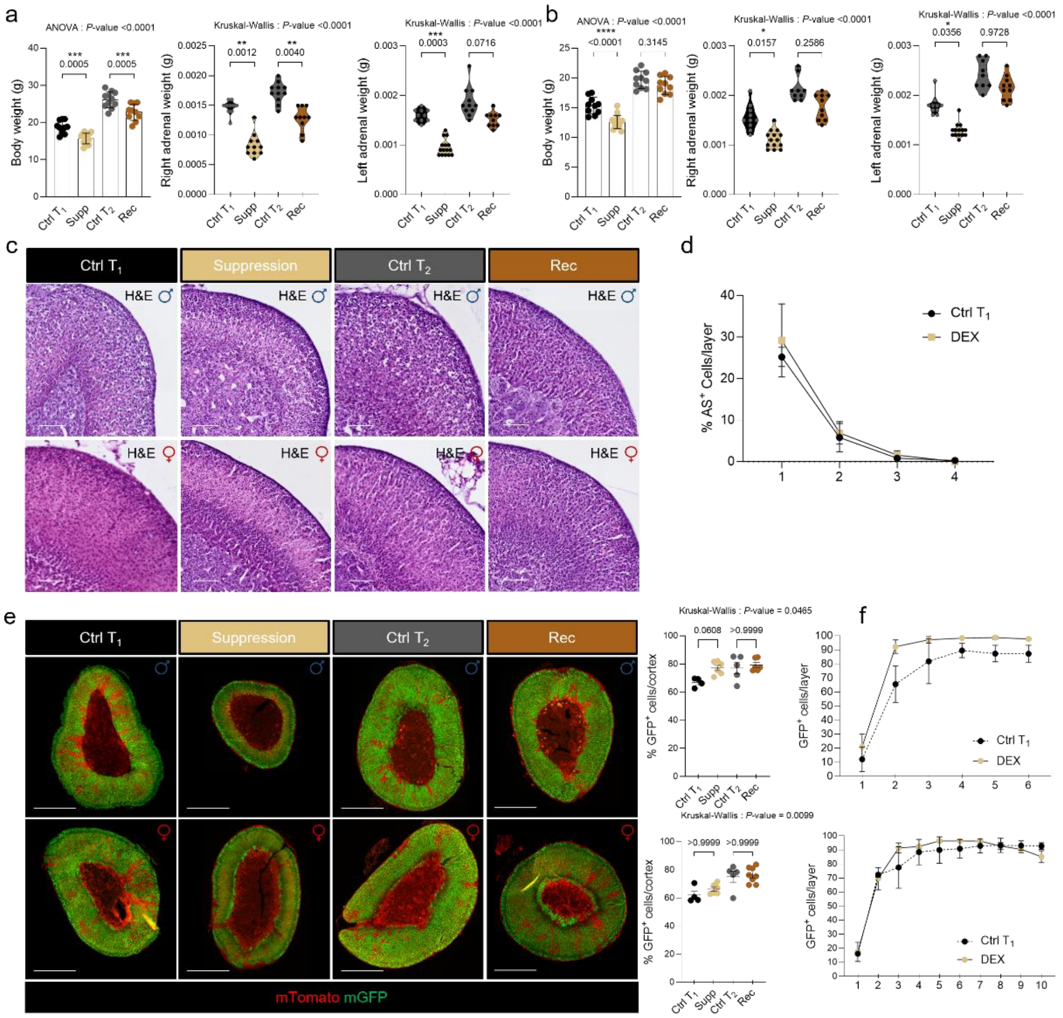


**Figure S13. Effect of dexamethasone treatment on the adrenal cortex.** Body weight and absolute right and left adrenal mass of *Cyp11b2*$^{+/Cre}$-*m*T*m*G male **(a)** and female **(a)** mice treated with dexamethasone (suppression) from 4 to 6 weeks of age ($T_1$) and after a recovery period from 6 weeks to 9 weeks ($T_2$). **(c).** H&E staining of male and female, control and treated, *Cyp11b2*$^{+/Cre}$-*m*T*m*G mouse adrenals at $T_1$ and $T_2$. **(d).** Quantification of AS$^+$ cells in the first 4 layers of the adrenal cortex of *Cyp11b2*$^{+/Cre}$-*m*T*m*G male mice treated with dexamethasone (DEX) from 4 to 6 weeks of age ($T_1$). Each layer has a thickness of 50 micron. **(e).** Coimmunofluorescence of mGFP (green) with mTomato (red) and quantification of the mGPF$^+$ cells in the adrenal cortex of *Cyp11b2*$^{+/Cre}$-*m*T*m*G (upper panel) male and (lower panel) female mice at $T_1$ and $T_2$. Scale Bar: 500 µm. **(f).** Analysis of the progression of mGFP+ cells along adrenal cortex layers of *Cyp11b2*$^{+/Cre}$-*m*T*m*G (upper panel, 6 layer) male and (lower pannel, 10 layer) female mice treated with DEX from 4 to 6 weeks of age ($T_1$) and after a recovery period from 6 weeks to 9 weeks ($T_2$). Each layer has a thickness of 25 micron; in males, only 6 layers could be quantified due to ZF atrophy. P values were obtained with Ordinary one-way ANOVA, adjusted with Šídák's multiple comparisons test or with Kruskal-Wallis test and adjusted with Dunn's multiple comparisons test, as indicated. Data are presented as mean values ±SEM. *P value < 0.05; **P value < 0.01; ***P value < 0.001.

**Figure S14**

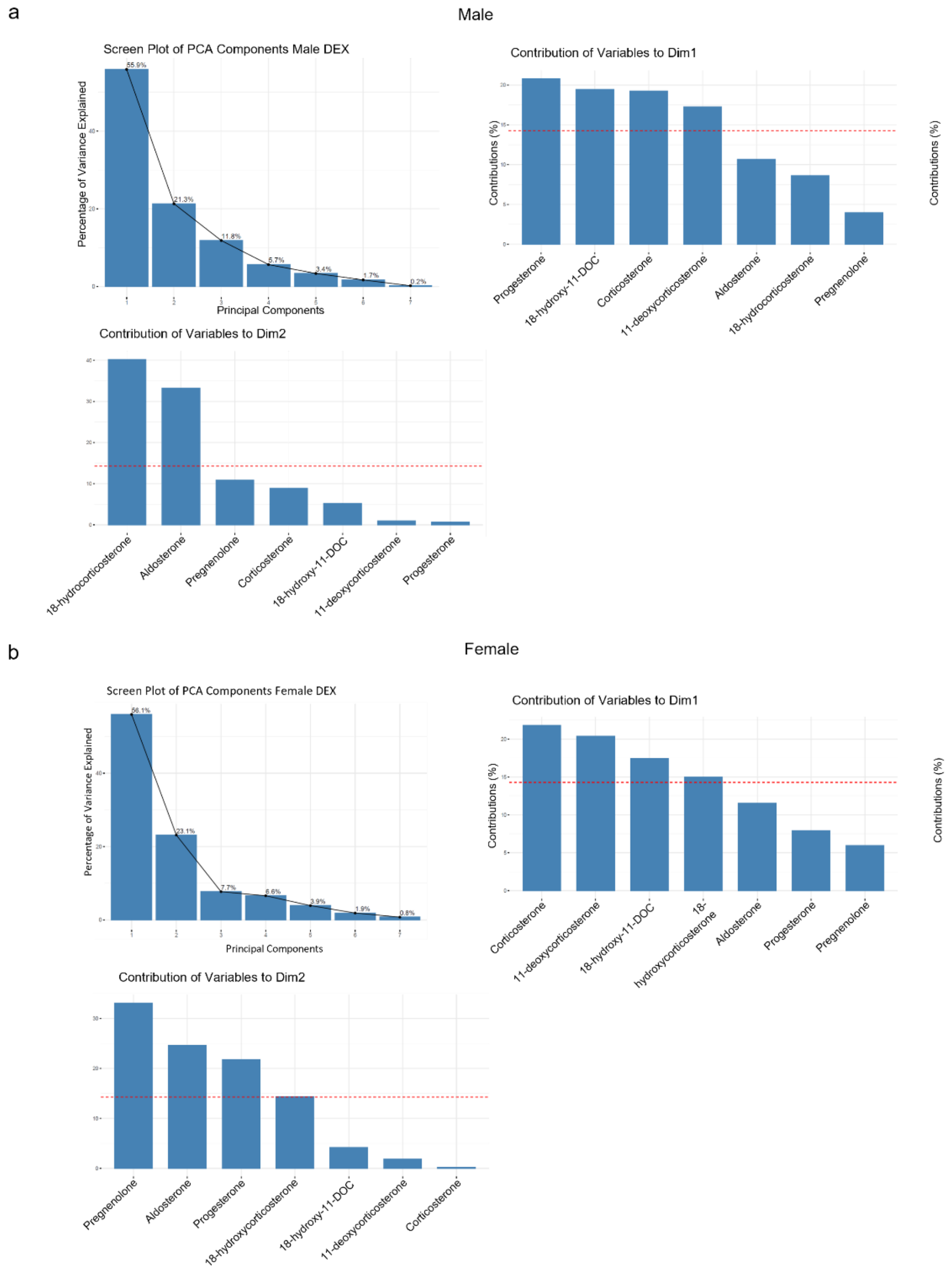


**Figure S14. Characteristics of Principal component analysis (PCA) of adrenal plasma steroid profiles following DXM treatment. (a-b).** Scree plot representing the principal components in relation to the percentage of explained variance and the respective contribution of single variables to dimensions 1 and 2 in male **(a)** and female **(b)** *Cyp11b2*$^{+/Cre}$-*m*T*m*G mice following DXM treatment vs control littermates. PCA shows clear separation between controls and DXM-treated mice along the first Dimension (Dim1), which explained 55.9% in males and 56.1% in females of the total variance, driven mainly by corticosterone, 11-deoxycorticosterone, 18-hydroxy-11-DOC, progesterone (only in males) and 18-hydrocorticosterone (only in females).

**Figure S15**

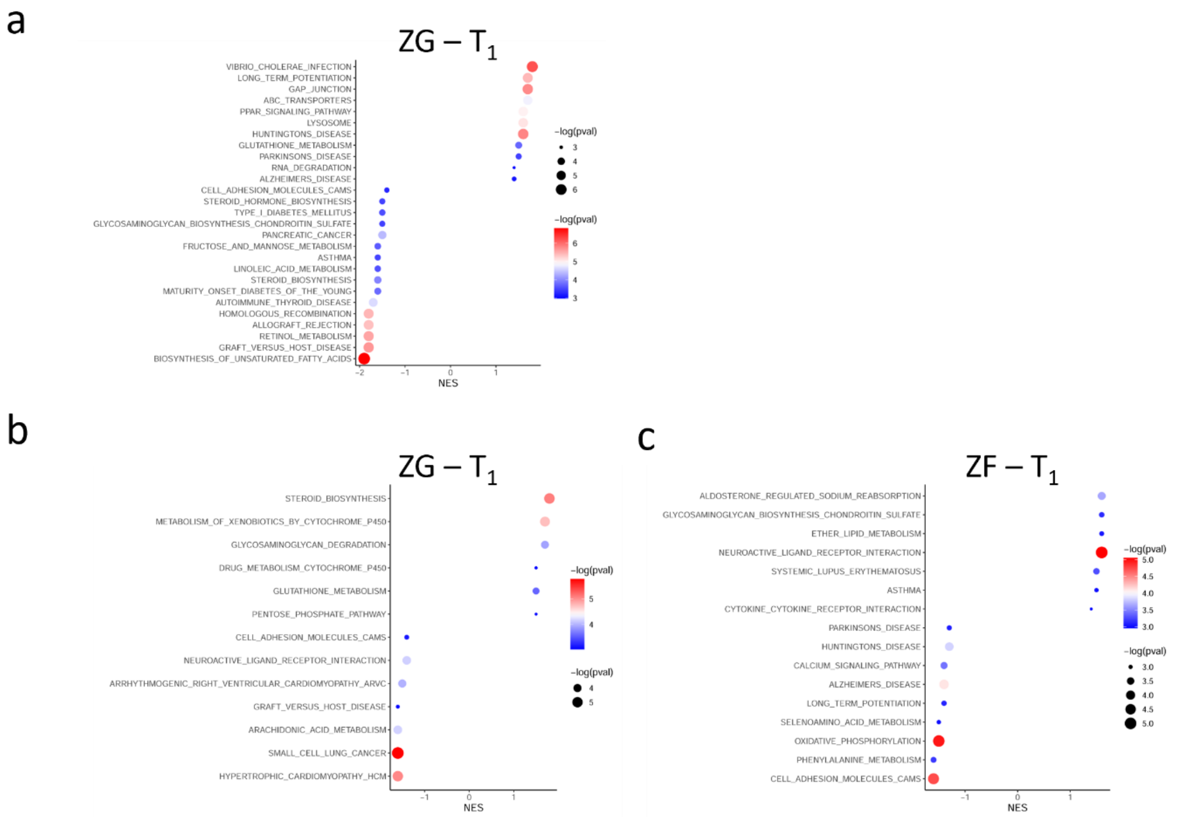


**Figure S15. Effect of DEX treatment on zone-specific gene expression (a-b).** Gene set enrichment analysis (GSEA) pathways from the KEGG database (adapted on mouse) of the ZG cluster of male **(a)** and female **(b)** *Cyp11b2*$^{+/Cre}$-*m*T*m*G DEX-treated vs Ctrl $T_1$ mice. **(c).** Gene set enrichment analysis (GSEA) pathways from the KEGG database (adapted on mouse) of female *Cyp11b2*$^{+/Cre}$-*m*T*m*G DEX-treated vs Ctrl $T_1$ mice in the ZF cluster.

**Figure S16**

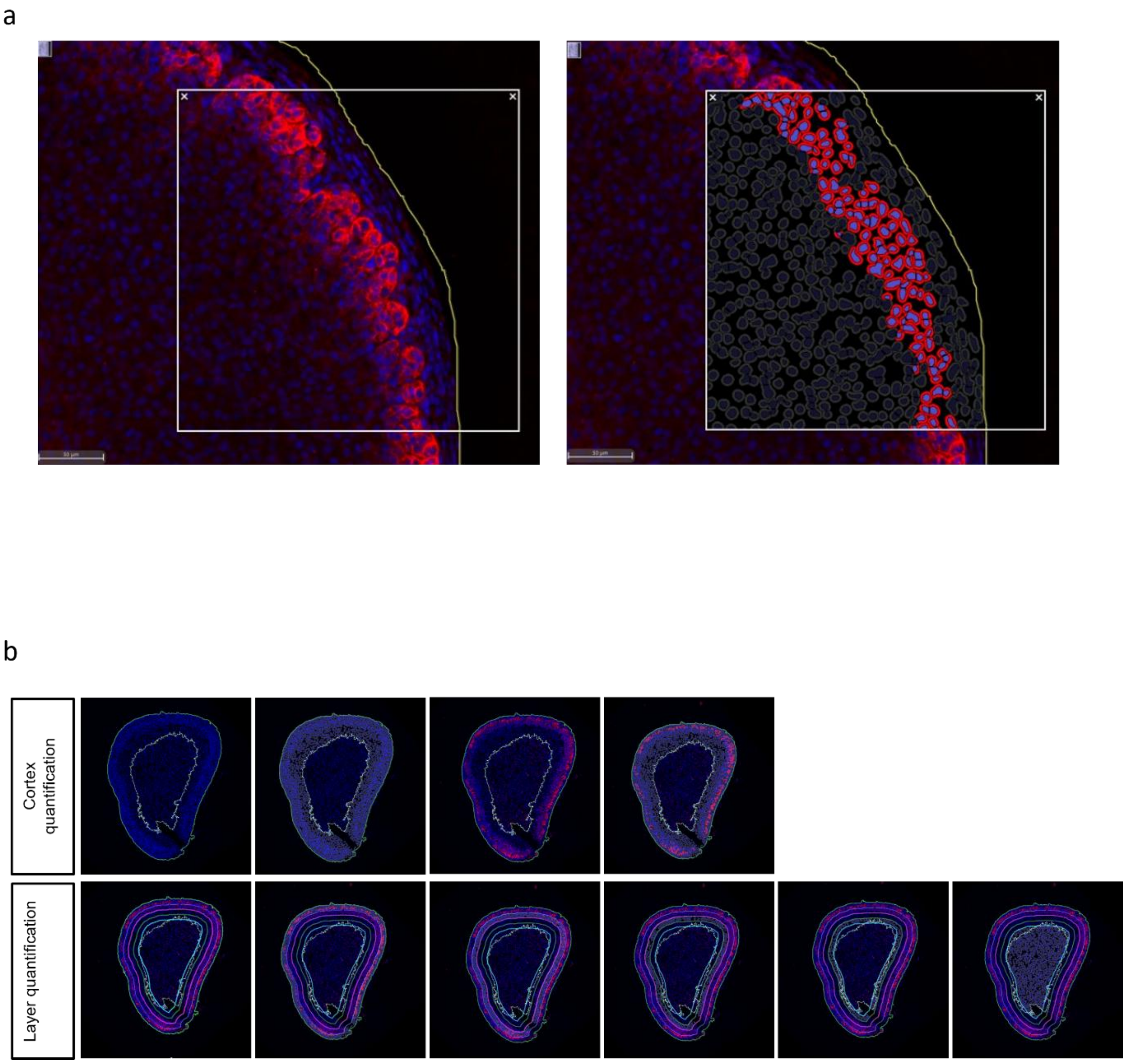


**Figure S16. Quantification of immunolabeling signals in the adrenal cortex (a).** Example of whole adrenal cortex DAB2$^+$ cells quantification using HALO software (version 3.6.4134, Indica Labs) with the HighPlex Fl v4.2.14 module. **(b).** Example of adrenal cortex segmentation in 50 µm layers of a DEX-treated male mouse and quantification of DAB2$^+$ cells using HALO software (version 3.6.4134, Indica Labs) with the HighPlex Fl v4.2.14 module.

### Figure S17

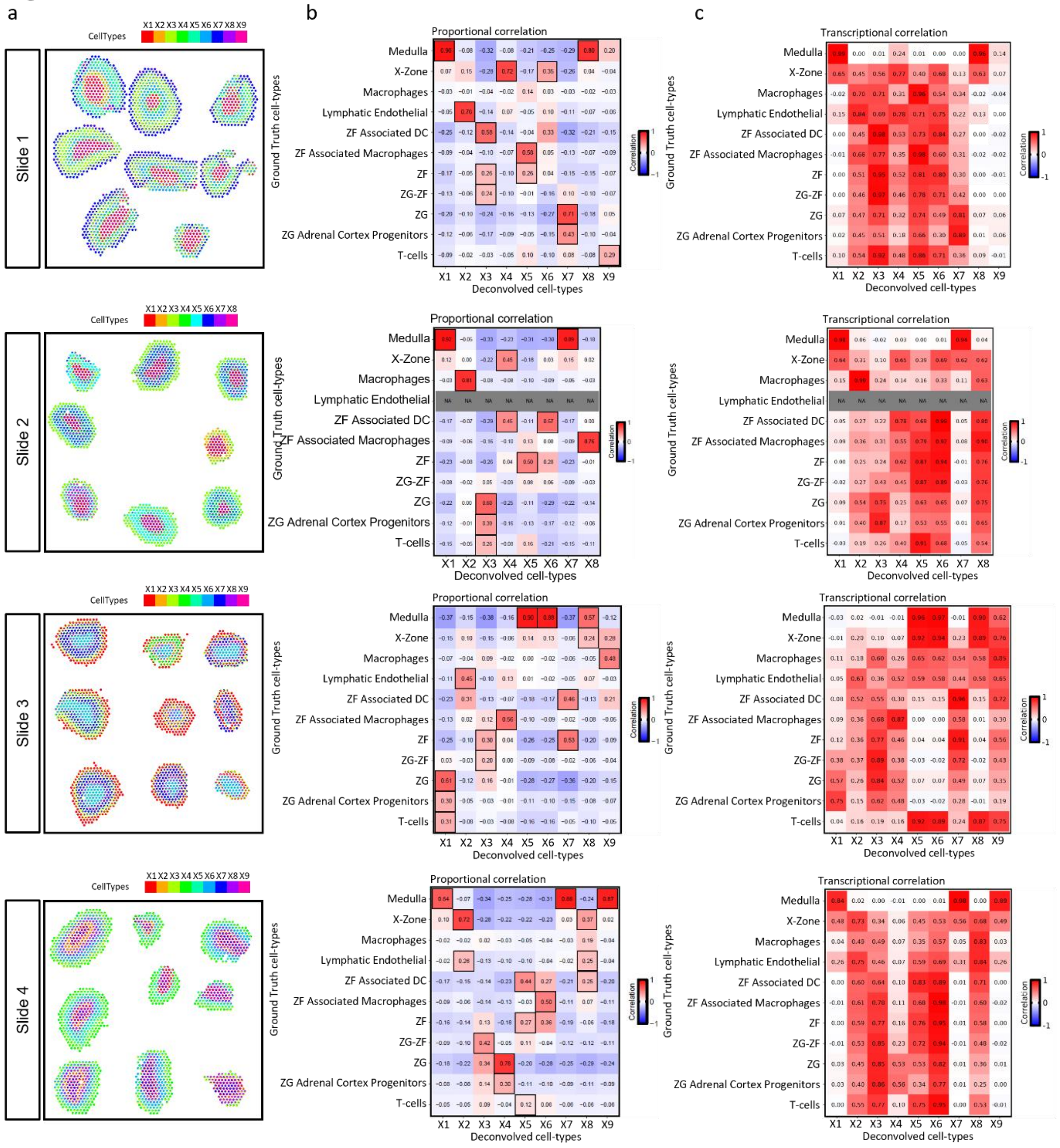


**Figure S17. Deconvolution of spatial transcriptomic data. (a).** Deconvolved cell-type proportions of adrenals ST data, displayed as pie charts for each ST spot. **(b).** Heatmap of Pearson's proportional correlations and transcriptional correlations between the deconvolved cell-type and ground truth cell-type proportions across spots. Matched deconvolved and ground truth cell types are boxed. **(c).** Heatmap of Pearson's transcriptional correlations between the deconvolved cell-type and ground truth cell-type proportions across spots.
